# Engineering Human ARMMs as Therapeutic Non-Viral Vehicles for *in vivo* Delivery of Genome Editing Payloads

**DOI:** 10.1101/2024.09.24.614725

**Authors:** Qiyu Wang, Shu-Lin Liu, Wen-Ning Zhao, Ryan von Kleeck, Michael Thomas, Kristin Luther, Zahra Vahramian, Yumei Sun, Komal Vyas, Pearl Moharil, Chris Chatzinakos, Carson O. Semple, Jennifer A. Badji, Leah Gens, Alysia Bryant, Steve Greenway, Chou-Wei Chang, Adam Francoeur, Nedyalka Valkov, Killian O’Brien, Joseph F. Nabhan

## Abstract

The ability to edit the human genome to treat previously incurable diseases or to modify previously undruggable targets has the potential to transform the lives of patients. However, delivery of genome editing machineries *in vivo*, and in a transient manner, continues to be a major challenge. We engineered human cell-derived ARRDC1-mediated microvesicles (ARMMs) as non-viral vehicles that mediate intracellular delivery of proteins, including DNA-modifying enzymes such as Cre recombinase and Cas9/guide RNA ribonucleoproteins (RNPs). ARMMs packaged multiple payloads per vesicle in our production system, yielding dose-dependent functional delivery of Cre or Cas9/gRNA RNP to a variety of cell types, including primary mouse and human cells. Oropharyngeal aspiration of ARMMs loaded with Cre led to biodistribution to alveolar macrophages (AMs), whereas intravenous administration predominantly to Kupffer cells (KCs), liver sinusoidal endothelial cells (LSECs), and splenocytes. Suprisingly, one administration of ARMMs loaded with Cas9 RNPs by oropharyngeal aspiration or intravenously resulted in efficient knockout of multiple genes in AMs or KCs, suggesting potential therapeutic utility. Through intravenous administration of ARMMs loaded with Cas9/NLRP3 gRNA we showed amelioration of drug-induced liver injury in a mouse model, supporting the use of ARMMs as human cell-derived therapeutic vehicles for genome editor delivery *in vivo*.

## Introduction

Recent progress in developing new delivery modalities has been instrumental in generating transformative therapies for patients with previously incurable disorders ^1^ ^2^ ^3^ and to combat infectious diseases ^3^. A number of these advances were made possible by delivery of therapeutic mRNA, RNA interference, or gene editing payloads. Conceptualization and realization of these therapeutic strategies has required a concerted effort from scientists across different disciplines to match therapeutic payloads with vehicles, viral or non-viral, to enable *ex vivo* or *in vivo* delivery to disease-relevant cell types of interest. However, significant hurdles continue to limit *in vivo* delivery of genome editors. These challenges are multi-dimensional and include physical constraints such as packaging efficiency and cargo capacity, safety concerns due to viral protein content, immunogenicity limiting redosability, toxicity ^4^ including off-target activity when delivering genome editors ^5–7^, and difficulties scaling production ^8,9^.

We developed and optimized a previously reported approach to engineer a class of cell membrane-derived extracellular vesicles, called ARRDC1-mediated microvesicles (ARMMs) ^10,11,12^, for *in vivo* delivery of a variety of protein payloads such as mCherry or Cre to monitor biodistribution, or therapeutic Cas/gRNA complexes otherwise referred to as RNPs. Engineered ARMMs are produced in human cells and are co-opted to intraluminally load the desired cargo through cellular expression of payloads directly tethered by a flexible peptide linker to ARRDC1. Concurrently, through overexpression in producer cells, ARMMs are decorated with a fusogen to ascertain maximal release of payload into the cytosolic compartment of recipient cells. For therapeutic applications, delivering a finite amount of RNPs is desirable as it may limit the possibility of off-target gene editing activity ^6,7^ and other reported genotoxicities ^5,13^ ^14^ ^15^. Additionally, using human-derived vesicles lacking a high viral protein load is less likely to elicit an immune response. Finally, scaling production is achievable in processes that resemble those used to generate therapeutic antibodies and other cell-derived biologics ^16^. While synthetic nanoparticles overcome some of these hurdles, dose-limiting toxicities, immunotoxicities, the inherent adjuvant-like properties and anti-drug antibodies have been reported ^17–19^. In this study, we show that ARMMs can intraluminally load and functionally deliver mCherry, Cre, and Cas9/gRNA RNPs to a range of cell types *in vitro* and *ex vivo*. Furthermore, we identify through oropharyngeal aspiration or intravenous administration of ARMMs, tissue resident macrophages, in the liver and lung, as cell types that have a strong propensity to internalize ARMMs and their corresponding payloads. Surprisingly, we discover that ARMMs have an exquisite ability to functionally deliver genome editing payloads *in vivo* to Kupffer cells in the liver, and to alveolar macrophages in the lung – and, in this context to efficiently target genes encoding both druggable and undruggable immune modulation proteins. *In vivo* Cas9 editing conveyed by ARMMs was shown to be efficient against multiple targets, including NLRP3 and IRF5, and able to provide protection against drug induced liver injury in a mouse model. Our data suggest that ARMMs could be used to genetically target such immune modulation nodes in macrophages to potentially produce therapeutic outcomes. This approach outlines a novel gene editing strategy to potentially treat complex diseases of the liver such as drug-induced liver injury.

## Results

### Packaging and functional delivery of payloads using engineered ARMMs

We first evaluated the loading of ARMMs with several proteins (mCherry, GFP, Cre) as cargo. This was achieved through expression of the desired payload as a single polypeptide chain fused to the ARRDC1 C-term, and separated by a flexible linker (**Fig.1A**). We investigated multiple approaches to package payloads in ARMMs. While previous work had relied predominantly on plasmid transfection-based loading of ARMMs ^10,11^, we explored using suspension and serum free-adapted HEK293-derived cells, called Expi293 (Thermo Fisher Scientific), as stable cell lines for production of protein-loaded ARMMs. Using a lentiviral transduction approach we generated a stable producer cell line of ARMMs loaded with GFP. To limit the potential impact of random integration of the transgene encoding the ARMM-loading cassette, we also developed an Expi293 cell line that harbors a Jump-In (Thermo Fisher Scientific) landing pad in the AAVS1 locus to introduce the ARMM loading transgene for mCherry. Consistent with previous reports ^11^, cellular production of engineered ARMMs loaded with Cre was achieved by plasmid transfection. Stable cell line production yielded ARMMs loaded with GFP or mCherry, suggesting that ARMMs production can be performed using either a transient transfection or a stable producer based approach (**Fig.1B**). The ability to potentially produce engineered ARMMs using stable suspension-adapted cell lines would more readily enable scaling production and reducing cost to support large scale manufacturing. Protein cargo loading in ARMMs was evaluated across three batches for each approach by immunoblotting using recombinant ARRDC1 for standard curve generation (**Fig.1B** and **Supplementary Fig.S1**). Fluorescent reporter proteins GFP or mCherry were loaded in ARMMs at an efficiency of >100 molecules/ARMM and Cre at an efficiency of ∼50 molecules/ARMM (**Supplementary Fig.S1**). Intraluminal payloads displayed immunopositive bands that correspond to the cargo, including ubiquitinated forms, as previously reported ^10^. Syntenin was used as an extracellular vesicle marker ^20^ and was enriched in purified ARMMs whereas vinculin was absent indicating no contamination with cells or cellular debris.

**Fig. 1.**
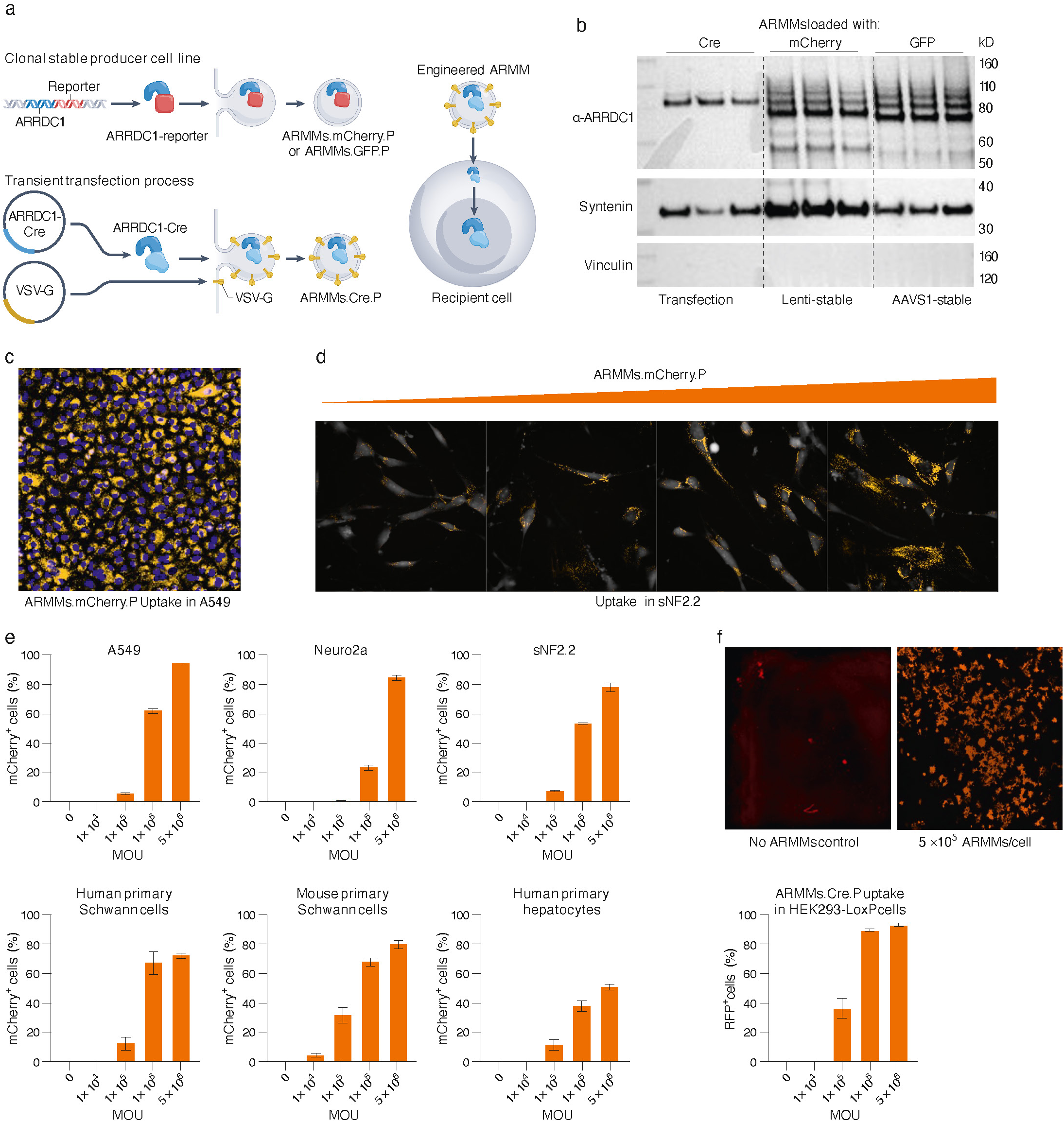
Production of engineered ARMMs and intracellular delivery of payloads. (A) Illustration of the generation of ARMMs loaded with different cargo proteins. Cargo proteins are expressed as fusion proteins to ARRDC1 via a flexible linker. Fusogens, such as VSV-G, are co-expressed to decorate budding ARMMs for improved functional delivery. Nuclear payloads included nuclear lead sequences to permit translocation to nucleus in recipient cells. (B) Representative Western blot analysis of engineered ARMMs loaded with Cre, mCherry or GFP. ARMMs were produced, as noted, by transient transfection of producer cells or from stable cell lines. Syntenin, an extracellular vesicle marker, is enriched in purified ARMMs and used as an EV loading control. Vinculin, a cellular marker, is absent in ARMMs, indicating no contamination with cell lysates or debris. Three batches of ARMMs are shown in the Western blot. (C) ARMMs.mCherry.P uptake in human lung alveolar epithelial cells A549. The representative image shows an intracellular mCherry signal (yellow) in a majority of cells (nuclear DAPI counterstain in blue). (D) Dose-dependent uptake of ARMMs.mCherry.P in a human schwannoma cell line sNF2.2, and (E) in other cell types, quantified by flow cytometry. (F) Functional delivery of nuclear protein Cre by ARMMs in a Cre-reporter cell line (HEK293T LoxP cells). Representative images show activation of tdTomato expression in the Cre reporter cell line upon treatment with ARMMs.Cre.P (24h). Cells treated with increasing MOU of ARMMs show dose-dependent effect by flow cytometry analysis.

Cargo delivery by ARMMs to recipient cells was first examined using live cell imaging. ARMMs loaded with GFP protein (ARMMs.GFP.P) were applied to A549 cells and their uptake was recorded over a 48h time period by fluorescence microscopy (**Supplementary videos**). The internalization of ARMMs.GFP.P was detectable within the first 2h and the accumulation of signal increased over 24h lasting through 48h when the experiment was terminated. Uptake of ARMMs was clear and dose-dependent. We quantified intracellular GFP signal delivered by ARMMs, and observed as puncta, using high-content imaging (**Supplementary Fig.S2**). ARMMs.GFP.P uptake was detected at a low MOU (multiplicity of uptake) of 1×10^4^ particles/cell and increased up to 5×10^7^ particles/cell, where saturation became visible, with no further increase in signal at higher MOUs. We expanded our evaluation of ARMMs uptake to a variety of cell types, including primary and post-mitotic cells (**Fig.1C, D, E**). To quantify uptake in vitro, cultured cells were treated with ARMMs loaded with mCherry (ARMMs.mCherry.P) for 24h and analyzed by flow cytometry (**Fig.1E**). All cells readily took up ARMMs.mCherry.P however uptake efficiencies varied across cell lines (**Fig.1E**), consistently attaining >50% transfection irrespective of cell type at higher MOU. These data support the notion that ARMMs can deliver payloads across a large range of cell types.

One of the major limitations of non-viral delivery platforms is the diminished ability to deliver payloads functionally due to trapping of payloads in sub-cytosolic compartments. We sought to determine if ARMMs loaded with Cre, decorated with the fusogen VSV-G and generated in our Expi293 suspension producer cells can mediate functional delivery of Cre. We used a Cre reporter cell line, HEK293-LoxP, in which the GFP expression cassette is flanked by LoxP sites followed by an RFP expression cassette (**Supplementary Fig.S3**). In the absence of Cre recombinase activity, GFP is expressed, whereas in the presence of Cre recombinase activity, the GFP expression cassette is excised and RFP is expressed. The transition from GFP to RFP in cells is indicative of functional Cre delivery. Reporter cells treated at an MOU of 1×10^6^ particles/cell showed ∼86% red fluorescence indicating that the Cre payload was delivered functionally using ARMMs to the nuclei of recipient cells (**Fig.1F**). Dose-dependent activation of RFP in treated cells was observed in ∼ 90% of cells at 1×10^6^ particles/cell.

We next treated primary cells isolated from Cre reporter mice, Ai9 and Ai14 ^21^, with ARMMs.Cre.P upon which functional delivery of Cre results in tdTomato expression. Lung macrophages, liver sinusoidal endothelial cells (LSEC), skin fibroblasts, and bone marrow derived macrophages (BMDM) were isolated from Cre reporter mice (**Fig.2A** and **Supplementary Fig.S4**). Primary cell cultures were subjected to treatment with ARMMs.Cre.P and monitored for tdTomato activation by fluorescence microscopy. 48 hours post-treatment, cells were fixed and stained for cell type-specific markers to confirm corresponding cell identity (**Fig.2B**). TdTomato expression was detected across all primary cell cultures and increased in a dose-dependent manner with higher MOU of ARMMs. Image analysis was performed to quantify cells expressing tdTomato (**Fig.2B**, right panel). In primary lung macrophages, at the lowest dose (1×10^4^ ARMMs/cell), ∼70% cells were tdTomato positive. These data indicate the high efficiency of functional delivery in this cell type. In LSECs, skin fibroblasts, and BMDM, 40-60% were detected at 1×10^5^ ARMMs/cell and 70-80% at 1×10^6^ ARMMs/cell. The dose-dependent effect indicates the successful functional delivery of Cre by ARMMs in primary cell types from various tissue types. The ability to functionally deliver intracellular payloads via ARMMs to macrophages with marked efficiency is noteworthy, and to our knowledge has not been reported by other non-viral vehicles. Collectively, these data indicate that cargo proteins can be efficiently packaged into ARMMs and delivered functionally across a very wide range of recipient cells.

**Fig. 2.**
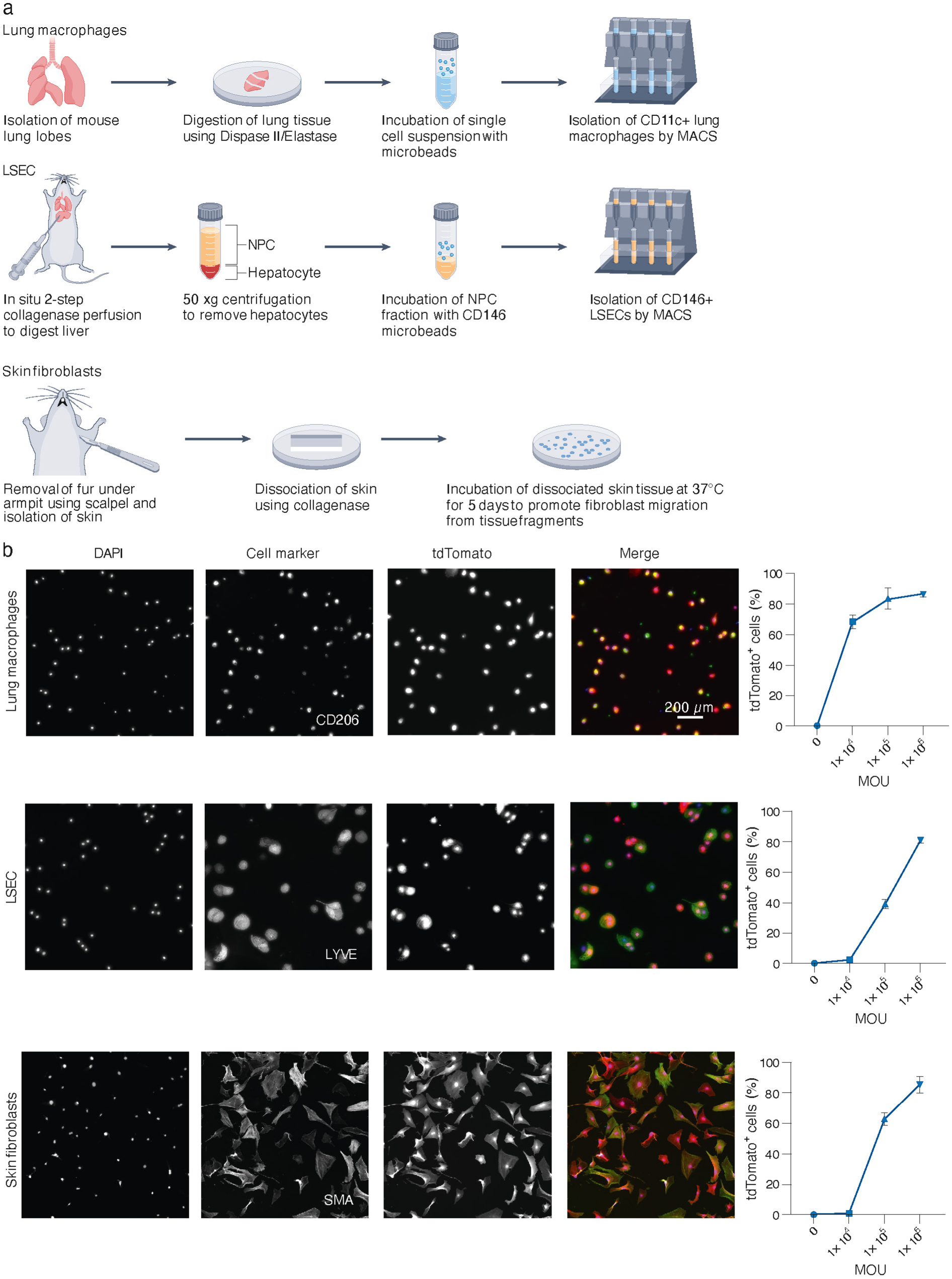
Functional uptake of engineered ARMMs in primary mouse cells. (A) Depiction of methodology used for isolation of three primary cell types from Ai14 Cre-reporter mice, including lung macrophages, liver sinusoidal endothelial cells, and skin fibroblasts. (B) Primary cells were incubated with ARMMs.Cre.P at increasing doses (MOU= 1×104, 1×105, or 1×106 ARMMs per cell) for 48h. Cells were fixed, stained with cell-type specific markers to confirm purity, and quantified for expression of tdTomato. Images display representative fields of view for DAPI (blue), appropriate cell marker (green), tdTomato (red), and merged channels for primary cells treated with 1×106 ARMMs per cell. Graphs displayed on the right indicate tdTomato positive cells out of total nuclei and error bars depict standard deviation from three replicates (>600 cells) per concentration.

### Engineering ARMMs as non-viral vehicles for delivery of CRISPR/Cas9 RNPs

Having demonstrated delivery of the nuclear protein Cre by ARMMs, we sought to explore if a similar approach could be used to functionally deliver Cas9/gRNA genome editing complexes across various cell types. We followed the same strategy detailed for other payloads to load Cas9 protein in ARMMs. To enable loading of fully functional holo-complex, we co-expressed the gRNA in producer cells and purified ARMMs loaded with Cas9/gRNA ribonucleoprotein (RNP, **Fig. 3A**). Multiple batches of ARMMs loaded with Cas9/gRNA (ARMMs.Cas9-gRNA.RNP) were characterized by immunoblotting and loading efficiency was calculated (**Fig.3B** and **Supplementary Fig.S5**). Despite Cas9 being a large 160KDa protein and we are loading the Cas9/gRNA RNP complex, ∼60-80 molecules of Cas9 were packaged per ARMM. To evaluate packaging of gRNA, we developed a universal quantitative PCR-based assay targeting a common scaffold sequence in Cas9 gRNA. Using standard curves generated with synthetic gRNA, the number of gRNA molecules loaded in ARMMs was calculated in the range of ∼40-120 copies/particle (**Supplementary Fig.S5**).

**Fig. 3.**
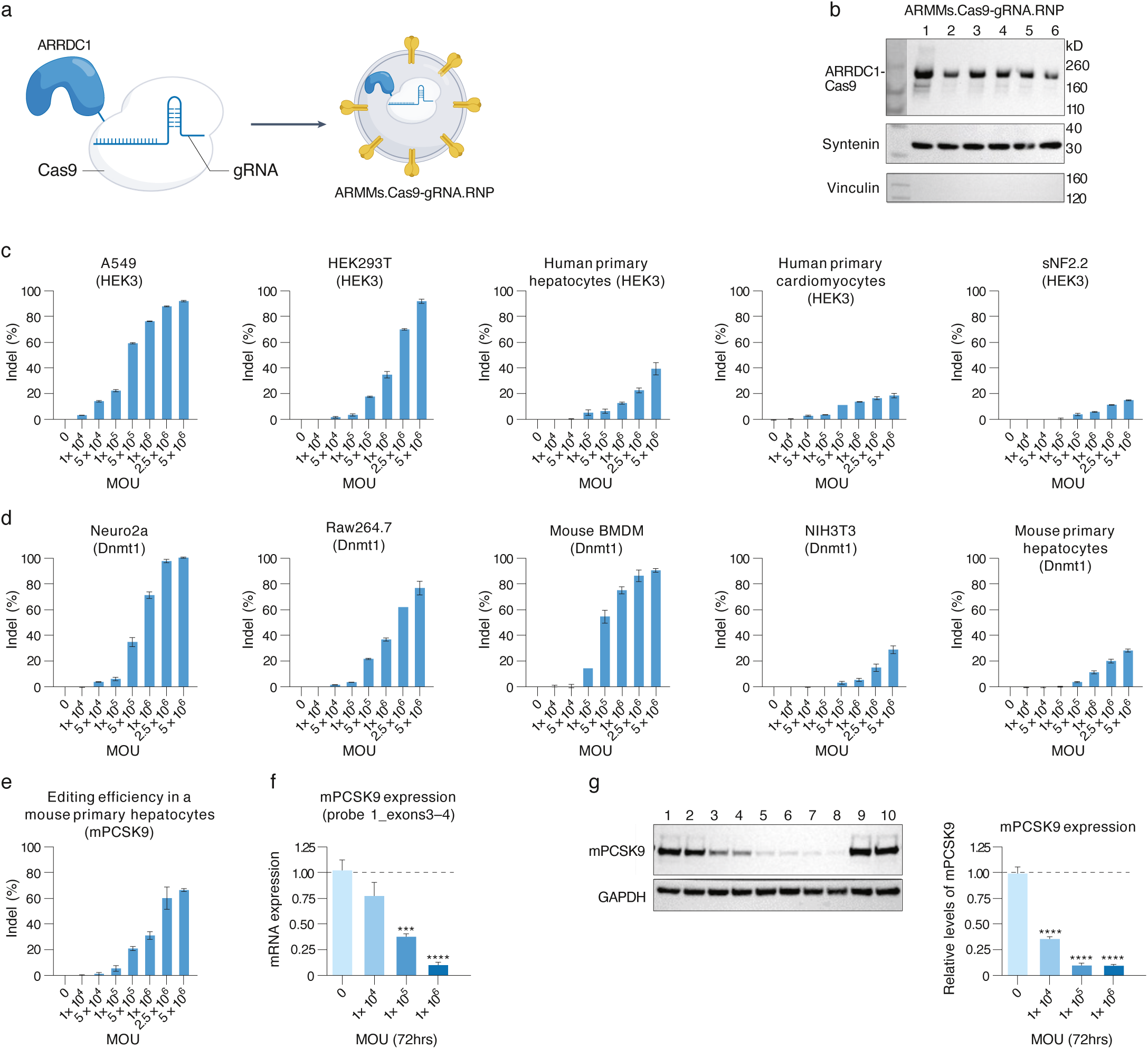
Functional delivery of ARMMs loaded with Cas9-gRNA RNPs in diverse cell types. (A) Illustration of approach to load ARMMs with Cas9/gRNA RNP. Cas9 protein is expressed as a fusion protein to ARRDC1 via a flexible linker. U6-driven co-expression of gRNA is derived from a separate plasmid to achieve loading of Cas9/gRNA RNP in engineered ARMMs. (B) Western blot analysis of ARMMs loaded with various Cas9-gRNA RNPs (5×10^9^/well) from different production batches. Syntenin, an extracellular vesicle marker, is enriched in purified ARMMs and is used as an EV gel loading control. Vinculin, a cellular marker, is absent in ARMMs, indicating no contamination with cell lysates. (C) Treatment of indicated cell types with increasing MOU of ARMMs loaded with Cas9 and a gRNA targeting the human HEK3 locus or (D) the mouse Dnmt1 gene yielded dose-dependent genome editing. Editing was quantified by NGS analysis and is shown as % indel. (E) Dose-dependent editing in mouse primary hepatocytes with ARMMs loaded with Cas9 and a gRNA targeting mouse Pcsk9 gene for 96h. (F) Quantitative PCR analysis showing reduced expression of Pcsk9 mRNA in primary hepatocytes upon treatment of ARMMs.Cas9-mPcsk9.RNP. (G) Western blot analysis showing reduced expression of PCSK9 protein upon treatment of ARMMs.Cas9-mPcsk9.RNP.

To evaluate functional delivery of genome editors by ARMMs, we chose two gRNAs that have been previously reported for high efficiency editing in a variety of cell types. Human HEK3 gRNA targets a highly accessible region in the human genome that does not encode any known genes ^22^ and mouse Dnmt1 gRNA targets the mouse Dnmt1 gene in exon 1 ^23^. We generated ARMMs loaded with Cas9 in complex with either HEK3 or Dnmt1 gRNA, and treated multiple cell types at increasing MOU of 1×10^4^ to 5×10^6^ particles/cell (**Fig.3C and D**). Cells were harvested after 72-96h and evaluated for gene editing at the target sequences by NGS. Editing efficiencies reported as % indel, as shown in **Fig.3C and D**, demonstrated dose-dependent editing in all cell types treated in this study. In A549, Neuro2a, and mouse bone marrow derived macrophages (BMDM), 70-80% indels were achieved at an MOU of 1×10^6^ particles/cell while ∼40% indels in HEK293T and Raw264.7 cells. Lower editing was detected in sNF2.2 and NIH3T3 cells. The lower editing efficiencies in the latter cell types cannot be completely attributed to reduced ARMMs uptake since, for example, at the treatment dose of 1×10^6^ particles/cell, ARMMs.mCherry.P uptake in sNF2.2 showed ∼55% mCherry^+^ cells (**Fig.1E**) whereas ∼5% indels were observed upon treatment with ARMMs.Cas9-HEK3.RNP. It is likely that the reduced editing efficiencies with the same gRNAs across different cell types is caused by diminished access to target genomic regions. To examine how ARMMs.Cas9-gRNA.RNP perform in post-mitotic cells, we treated primary human cardiomyocytes and hepatocytes or mouse hepatocytes with the ARMMs loaded with Cas9 and either HEK3 or DNMT1 gRNAs (**Fig.3C and D**). Editing was detectable at 5×10^5^ ARMMs/cell with a dose-dependent increase, though the overall editing efficiency was lower than other cell types.

To explore the potential of ARMMs-mediated delivery of Cas9 to edit a therapeutic target, we loaded ARMMs with Cas9 and a gRNA targeting exon 1 of mouse Pcsk9 (**Supplementary Fig.S6)** ^23,24^. We generated ARMMs.Cas9-mPcsk9.RNP and treated mouse primary hepatocytes at increasing MOU of 1×10^4^ to 5×10^6^ particles/cell for 72h. **Fig.3E** shows that editing is detectable at a low MOU of 5×10^4^ particles/cell and increases at higher doses. Next, we examined if Pcsk9 editing yielded reduced expression of mPcsk9 gene and protein. Treatment with ARMMs.Cas9-mPcsk9.RNP reduced mPcsk9 gene expression (**Fig.3F** and **Supplementary Fig.S6**). A dose-dependent decrease in PCSK9 was observed with ∼50-60% reduction at 1×10^5^ particles/cell treatment and 80-90% at 1×10^6^ ARMMs/cell. This translated into a dose-dependent reduction in mPcsk9 protein levels (**Fig.3G)**, with 35% or 10% residual protein left at 1×10^5^ or 1×10^6^ particles/cell treatments, respectively. Taken together, these data demonstrate that engineered ARMMs are efficient delivery vehicles for Cas9/gRNA RNP across diverse cell types.

Non-viral delivery of Cas9 mRNA and gRNA using lipid nanoparticles has been shown to induce innate immunity through activation of Toll-like receptor signaling ^25^. We sought to examine whether ARMMs inherently or when actively loaded with Cas/gRNA RNPs induce immune activation of Toll-like receptors (TLRs). We generated ARMMs loaded with mCherry protein or Cas9/gRNA RNP and treated mouse bone marrow derived macrophages (**Supplementary Fig.S7**). We posited that protein-loaded ARMMs would reflect activation by passively incorporated cell-derived nucleic acids, and RNP-loaded ARMMs due to actively incorporated gRNA. TNFα and IL6 in the cell culture supernatants were evaluated 6h after treatment and were not found to be increased by treatment with ARMMs indicating absence of TLR activation. These data support the ability of engineered human ARMMs to efficiently deliver Cas/gRNA RNP without inducing immunotoxicity.

### *In vivo* delivery of payloads by engineered ARMMs is efficient in mice

We first administered ARMMs.mCherry.P *in vivo* by oropharyngeal aspiration in C57BL/6 mice (Charles River Laboratories) to evaluate biodistribution in the lungs. 2 to 48 h later, tissues were harvested, processed and analyzed by ELISA, flow cytometry and immunohistochemistry (**Supplementary Fig.S8**). Cellular uptake of ARMMs was exclusively detected in alveolar macrophages, identified as CD45+CD11bmidCD11C+SiglecF+ (**Supplementary Fig.S9**). Delivery of the fluorescent protein payload mCherry via ARMMs.mCherry.P showed strong uptake in the alveolar macrophage population 2h after oropharyngeal aspiration as monitored by flow cytometry (**Supplementary Fig.S8B**), with ∼50-75% of alveolar macrophages displaying positive mCherry signal (**Supplementary Fig.S8C**). Histological assessment of paraffin embedded lung cross-sections showed strong mCherry antibody staining in CD206^+^ alveolar macrophages (**Supplementary Fig.S8D**). Furthermore, mCherry protein remained detectable in alveolar macrophages by histology 48h after administration (**Supplementary Fig.S8E**).

To investigate the potential for *in vivo* functional payload delivery using ARMMs, we evaluated local administration of ARMMs.Cre.P to the lung. A single dose of 5×10^11^ ARMMs.Cre.P was administered via oropharyngeal aspiration to Cre-reporter mice (**Fig.4A**). Four days after ARMMs.Cre.P aministration, lungs were harvested and tdTomato signal was assessed in alveolar macrophages. Alveolar macrophages isolated from Cre-reporter mice revealed a robust tdTomato signal in ARMMs.Cre.P-treated animals as determined by flow cytometry (**Fig.4B**), in comparison with PBS-treated control mice. ∼50-80% of isolated alveolar macrophages displayed tdTomato signal after one dose (**Fig.4C**). Robust functional delivery of Cre by ARMMs was detected across all lung lobes, with stronger tdTomato signals observed in the upper lobes (**Fig.4D**). Further analysis of tdTomato biodistribution by histology in fixed-frozen lung sections revealed strong colocalization of tdTomato signal with lung macrophage marker CD206 in the ARMMs.Cre.P-treated mice (**Fig.4E**), with no tdTomato signal detected in PBS-treated controls.

**Fig. 4.**
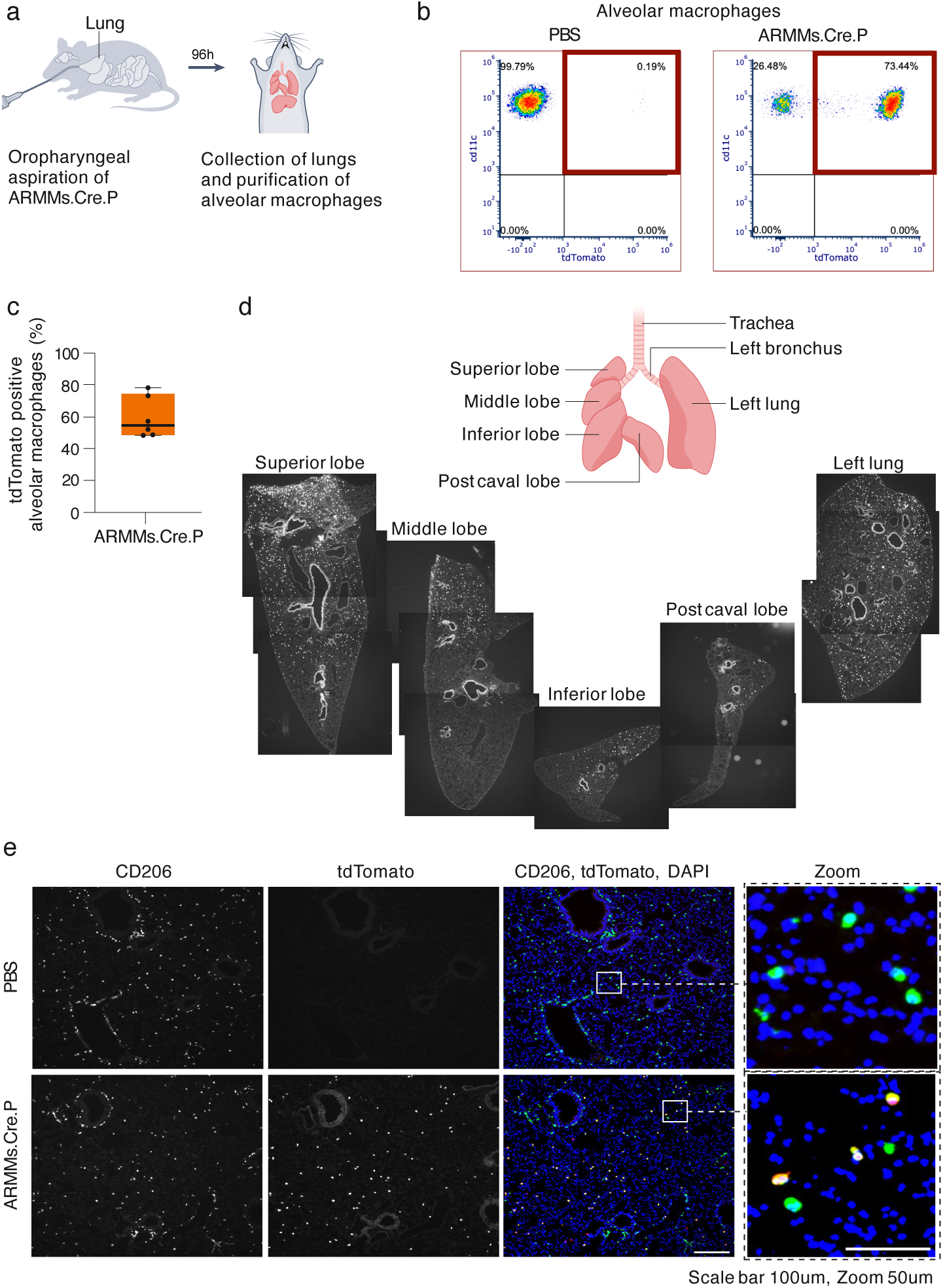
Oropharyngeal aspiration of engineered ARMMs results in functional delivery of payload to alveolar macrophages. (A) Schematic of in vivo dosing by oropharyngeal aspiration of engineered ARMMs with Cre recombinase in reporter mice. Lungs were harvested 96h after administration. (B) Representative flow cytometry density plots of tdTomato+ alveolar macrophages (see Supplementary Fig.S9 for gating strategy) from PBS or ARMMs.Cre.P-treated animals. (C) Percentage of tdTomato+ alveolar macrophages of total alveolar macrophages as determined by flow cytometry from mice (n=6) administered ARMMs.Cre.P. (D) Tissue cross sections of lung lobes from mice treated with ARMMs.Cre.P displaying tdTomato fluorescence. Functional delivery of ARMMs was observed across all lung lobes with higher tdTomato abundance observed in upper lobes. (E) Fixed frozen lung sections were stained for CD206 (macrophage marker) to visualize colocalization with tdTomato expressing cells.

We then evaluated tissue-specific biodistribution of ARMMs by intravenous administration. We performed a pharmacokinetic evaluation of ARMMs administered via retro-orbital intravenous (IV) injection of ARMMs.GFP.P (**Supplementary Fig.S10A**). GFP payload signal was measured in plasma collected from mice at several timepoints post-administration. No signal was detected 2h post-administration suggesting rapid biodistribution to target tissues and/or clearance. To survey tissue types for uptake of ARMMs 24h post-administration, tissue homogenates from PBS-perfused animals were analyzed by ELISA (**Supplementary Fig.S10B**). Several tissues showed a robust signal corresponding to the delivered payload, GFP, by ELISA. Uptake was detected in spleen, liver, kidney and lung, with the latter yielding lower signals. Importantly, repeat administration of ARMMs loaded with GFP protein in immunocompetent mice did not lead to generation of neutralizing antibodies that inhibit cellular uptake of ARMMs suggesting the possibility of repeat administration (**Supplementary Fig.S11**).

To investigate functional payload delivery by intravenously ARMMs administered to specific cell types, we injected Cre reporter mice with ARMMs loaded with Cre recombinase (ARMMs.Cre.P). A single dose of 1×10^12^ ARMMs.Cre.P was administered to Ai14 mice, four days after which, functional delivery of Cre to liver tissue was assessed by immunofluorescence of fixed tissue cross-sections and by flow cytometry of liver cell suspensions to evaluate tdTomato+ cells in various cell populations (**Fig.5 and Supplementary Fig.S12**). TdTomato colocalized strongly with Kupffer cell marker IBA1 (**Fig.5A**) and liver sinusoidal endothelial cell marker LYVE-1 (**Fig.5B**) in liver cross sections by immunofluorescence staining. Characterization of liver biodistribution by flow cytometry revealed that approximately 40-60% of the Kupffer cell population (CD45+F4/80+CD11bint) (**Fig.5C-D**) and 10-20% of the LSEC population (CD45-CD31+CD146+) (**Fig.5E-F**) were tdTomato positive.

**Fig. 5.**
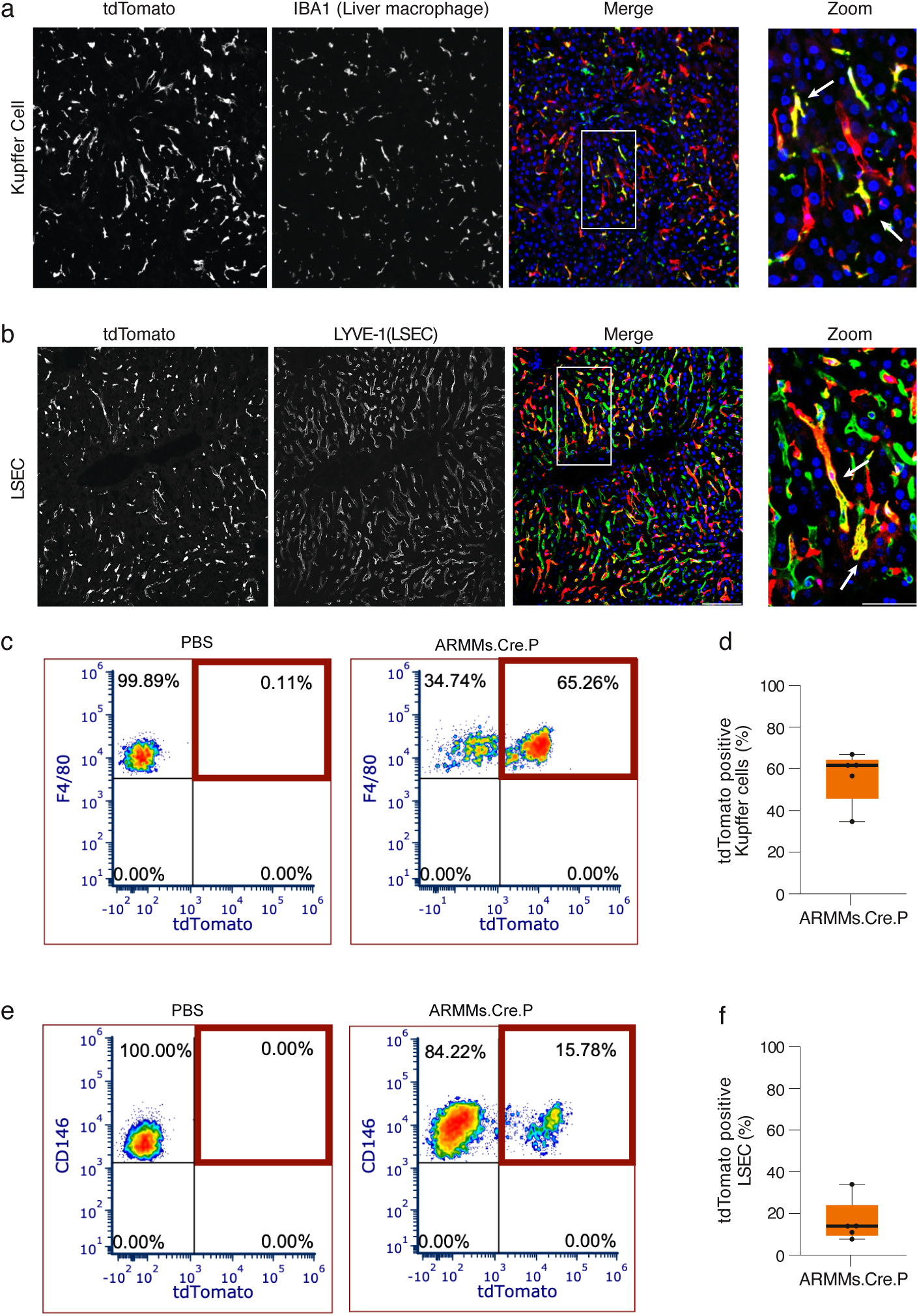
Intravenous administration of engineered ARMMs results in functional delivery of payload to liver Kupffer cells and liver sinusoidal endothelial cells (LSECs). Ai14 reporter mice received a single dose of 1×1012 ARMMs.Cre.P through retro-orbital injection. Functional delivery of Cre payload was evaluated after 4 days in the liver by histological and flow cytometry assessment of tdTomato expression in liver-derived cells. (A) Liver cross-sections stained for IBA1 (green, liver macrophage marker) and tdTomato. (B) Liver cross sections stained for LYVE-1 (green, LSEC marker) and tdTomato (native fluorescence). White arrows in “Zoom” images depict colocalization events (yellow). Single cell suspensions of non-parenchymal cells (NPCs) were evaluated for tdTomato expression by flow cytometry in (C, D) the Kupffer Cell (CD45+F4/80+CD11bint) population (n=5 mice) and (E,F) the LSEC (CD45-CD31+CD146+) population (n=5 mice). See Supplementary Fig.S12 for gating strategy.

### Engineering ARMMs as non-viral vehicles for genetic disruption of therapeutic targets

To investigate their therapeutic utility *in vivo*, we loaded ARMMs with Cas9 RNPs targeting mouse Nlrp3 or Irf5. The NLRP3 protein is a key component of the NLRP3 inflammasome, a critical mediator of immune responses in both sterile and infection-mediated inflammation ^26^. Similarly, IRF5 activation and involvement has been shown in multiple diseases of inflammation, including systemic lupus erythrmatosus and Sjogren’s ^27^.

To disrupt expression of Nlrp3 using Cas9, 19 gRNAs were screened by transient transfection in mouse Neuro2a cells (**Supplementary Fig.S13A**). Two gRNAs displayed the highest level of editing: gRNA #7 targeting the promoter region, and gRNA #11, targeting the NLRP3 coding region (**Supplementary Fig.S13B**).

We focused our effort on Nlrp3 gRNA #11 that showed the highest level of Nlrp3 coding sequence disruption. ARMMs loaded with Cas9 and NLRP3 gRNA #11 were generated (ARMMs.Cas9-Nlrp3.RNP). Editing efficiency by ARMMs.Cas9-Nlrp3.RNP was first evaluated in mouse RAW264.7 cells (**Fig.6A**) yielding ∼90% indels at 1×10^6^ MOU. The disruption of the Nlrp3 gene translated into significant reduction in NLRP3 protein levels in RAW264.7 cells and mouse bone marrow derived macrophages (**Fig.6B and Supplementary Fig.S14**). Treatment with LPS and Nigericin (NIG) is commonly used to stimulate NLRP3 inflammasome activity, leading to increased intracellular NLRP3 protein levels and pro-inflammatory IL-1β cytokine release. NLRP3 inflammasome stimulation by LPS and NIG was almost completely abolished in cells treated with ARMMs.Cas9-Nlrp3.RNP leading to loss of NLRP3 protein (**Fig.6B**) and blunting of IL-1β secretion (**Fig.6C**).

**Fig. 6.**
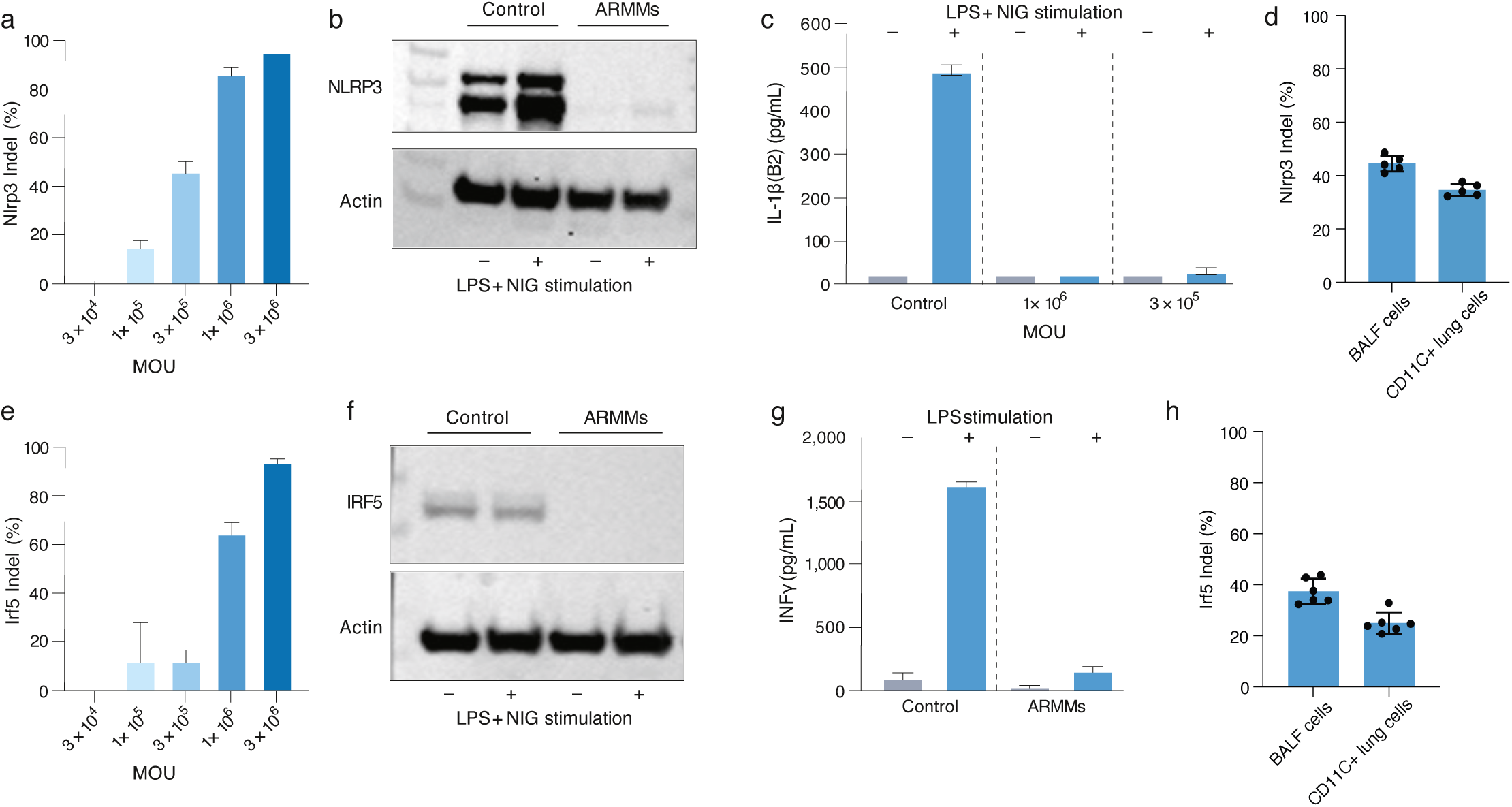
Delivery of Cas9-gRNA RNPs targeting NLRP3 or IRF5 by engineered ARMMs blunts pro-inflammatory cytokine production. (A) NGS analysis of genomic DNA extracted from macrophage cell line RAW264.7 after treatment with ARMMs.Cas9-Nlrp3.RNP showing dose-dependent gene editing. (B) Western blot analysis showing reduction of NLRP3 protein after treatment of RAW264.7 cells with ARMMs.Cas9-Nlrp3.RNP. LPS + Nigericin (NIG) treatment did not induce NLRP3 expression after treatment with ARMMs. (C) Anti-IL-1β ELISA of supernatants from BMDM ELISA after treatment with ARMMs.Cas9-Nlrp3.RNP or PBS followed by LPS + NIG. ∼20-fold induction of IL-1β was detected in mock-treated control cells. (D) In vivo administration of ARMMs.Cas9-Nlrp3.RNP by oropharyngeal aspiration results in editing of ∼30-40% of alveolar macrophages isolated from BALF (bronchoalveolar lavage fluid) and CD11C+ cells from lung tissue. Editing was evaluated by NGS analysis of genomic DNA. (E) NGS data analysis showing dose-dependent gene editing with ARMMs.Cas9-Irf5.RNP in RAW264.7 cells. (F) Western blot analysis showing reduction of IRF5 protein expression after ARMMs.Cas9-Irf5.RNP treatment. (G) anti-INFγ ELISA of supernatants from BMDM after treatment with ARMMs.Cas9-Irf5.RNP followed by LPS stimulation. ∼160-fold induction was detected in mock-treated control cells whereas <4-fold was detected in ARMMs-treated cells. (H) In vivo administration of ARMMs.Cas9-Irf5.RNP by oropharyngeal aspiration results in editing of ∼30-40% of alveolar macrophages isolated from BALF (bronchoalveolar lavage fluid) and CD11C+ cells from lung tissue. Editing was evaluated by NGS analysis of genomic DNA.

We extended the evaluation to a more challenging target to inhibit by small molecule approaches, the transcription factor IRF5. Identifying novel approaches to disrupt IRF5 expression is therefore desirable. We generated ARMMs loaded with Cas9 and an Irf5-targeting gRNA (ARMMs.Cas9-Irf5.RNP) and evaluated their activity in RAW264.7 cells. Dose-dependent editing was observed yielding ∼90% indels at 1×10^6^ MOU (**Fig.6E**). IRF5 expression was not detectable after treatment with ARMMs (**Fig.6F**) indicating robust knockout of Irf5, leading to blockade of IFNβ release after LPS stimulation (**Fig.6G**).

Since oropharyngeal aspiration of ARMMs.Cre.P showed robust functional delivery of nuclear protein Cre to alveolar macrophages (**Fig.4**), we resorted to the same route of administration to evaluate functional delivery of Cas9-gRNA RNP and efficiency of *in vivo* editing of Nlrp3 or Irf5. ARMMs loaded with Cas9-Nlrp3 or Cas9-Irf5 RNPs were administered via oropharyngeal aspiration. 96h later, cells were harvested from bronchoalveolar lavage fluid (BALF) or disgested liver tissue and macrophages were isolated and evaluated for editing of the Nlrp3 target region or the Irf5 target region (**Fig.6G**). A remarkably high efficiency of 40-50% indels for Nlrp3 and 30-40% for Irf5 were observed in these cells. Our data indicate that ARMMs are efficient non-viral vehicles for *in vivo* delivery of CRISPR-Cas9 genome editors to alveolar macrophages.

### ARMMs loaded with NLRP3-targeted Cas9 RNP ameliorate drug-induced liver injury

We extended our *in vivo* evaluation of Cas9 RNP delivery to intravenous administration of ARMMs. A single intravenous administration of ARMMs loaded with Cas9/NLRP3 gRNA yielded a high level of editing (∼60% indels) in Kupffer cells (**Fig.7A**). As mouse liver resident KC display a robust ability for *in vivo* genomic editing via ARMMs, we sought to evaluate ARMMs in a carbon tetrachloride (CCl4)-elicited mouse model of acute liver injury ^28^. Mice were treated with prophylactic IV injection of ARMMs.Cas9-Nlrp3.RNP 3 days prior to CCl4-elicited injury. Although pathology assessment of H&E stained liver sections did not show significant improvement in ARMMs.Cas9-Nlrp3.RNP treated mice (**Supplementary Fig.S15**), analysis of mouse serum collected 24 hours post-CCl4 administration demonstrated significant reduction (∼50%) in ALT and AST enzymes in ARMMs.Cas9-Nlrp3.RNP injected animals relative to vehicle control (**Fig.7B,C**). Furthermore, liver single cell resuspensions processed in flow cytometry indicated that CCl4 treatment produced a significantly diminished impact on cell viability in mice treated with ARMMs.Cas9-Nlrp3.RNP compared to vehicle control (**Fig.7D**) with about 2-fold increase in % of live cells compared to CCl4 treated controls. Additionally, ARMMs.Cas9-Nlrp3.RNP treatment resulted in noticeably diminished necrosis in liver cells (**Fig.7E**), though the scoring did not reach statistical significance. Congruently, tissue sections analyzed by TUNEL staining, reflective of apoptosis, showed ∼50% reduction in mice treated with ARMMs.Cas9-Nlrp3.RNP relative to vehicle control (**Fig.7F**). Altogether, these results illustrate the potential utility of ARMMs in targeting tissue resident Kupffer cells to ameliorate drug-induced liver injury.

**Fig. 7.**
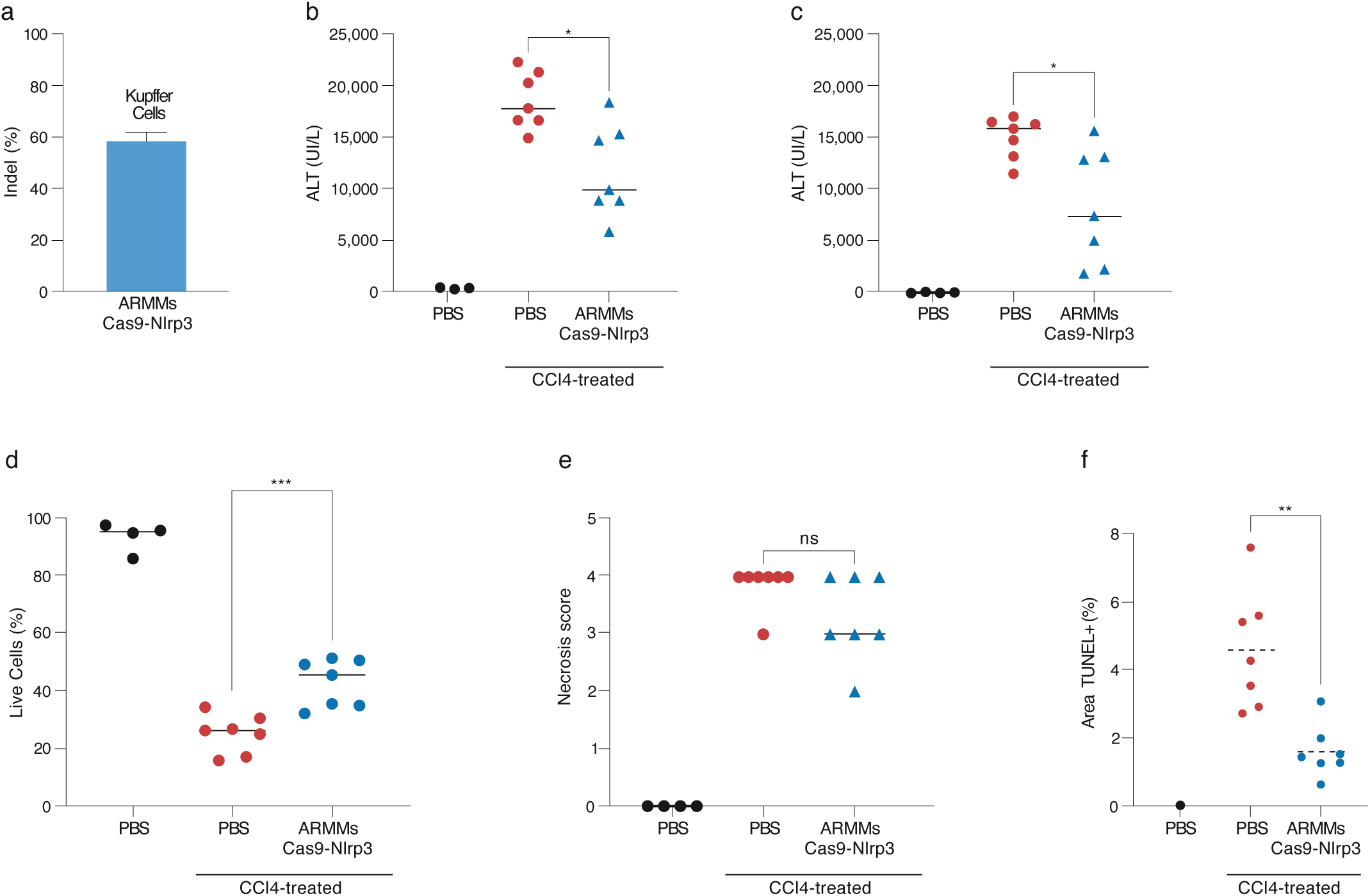
In vivo editing of NLRP3 by engineered ARMMs ameliorates acute liver injury. (A) Assessment of in vivo editing efficiency by NGS analysis of genomic DNA from isolated mouse liver Kupffer cells 3 days post-intravenous administration of ARMMs.Cas9-Nlrp3.RNP. (B-F) Mice were prophylactically treated with PBS vehicle control or ARMMs.Cas9-Nlrp3.RNP prior to inducing liver injury with CCl4. 24h post-treatment with CCl4, liver enzymes (B) ALT and (C) AST were assessed. (D) Percentage of live cells was determined by flow cytometry of liver single cell suspensions. (E) Necrosis score by H&E staining of liver tissue sections, and (F) percentage of TUNEL+ staining by immunofluorescence microscopy of liver tissue sections. Significance was determined relative to PBS control, CCl4 treated mice in one-way ANOVA, Dunnett’s multiple comparisons test. *p<0.05, **p<0.001, ***p<0.0001.

## Discussion

This work establishes ARMMs as a human cell-derived system that can be co-opted and engineered to treat complex and genetic diseases *in vivo*. Incorporation of protein or RNP payloads in ARMMs can be achieved through association via a flexible peptide linker with ARRDC1, which localizes to the inner leaflet of the plasma membrane during budding ^10^. Using this strategy, we demonstrated loading of ARMMs with reporter proteins mCherry and GFP, Cre recombinase, and Cas9/gRNA RNP using either stable producer cells or by transfection of plasmids encoding ARRDC1, the associated payloads and a fusogen. The ability to potentially use a stable producer cell line, grown in suspension and in a defined serum-lacking medium, to generate intraluminally loaded particles is an important step forward in enabling scalable production of ARMMs to support clinical trials ^16^.

The high efficiency of loading of engineered ARMMs yields potent non-viral vehicles that allow for robust delivery to recipient cells. ARMMs were readily internalized by a range of cells, including cancer cell lines such as A549 and Neuro2a, as well as primary neuronal cells or hepatocytes, suggesting that delivery is potentially achievable to any cell. Importantly, that ARRDC1 is not an integral membrane protein does not impede the ability to functionally deliver nuclear payloads such as Cre, which was demonstrated in several primary mouse cells including lung macrophages, LSECs, bone marrow-derived macrophages and skin fibroblasts. Higher doses of ARMMs produced high levels of functional delivery of payloads to treated cells further re-enforcing the notion that ARMMs can potentially mediate unrestricted delivery to any cell type. To realize the therapeutic potential of ARMMs, we demonstrated efficient loading with Cas9/gRNA RNPs at a concentration >60 RNPs/vesicle. Evaluation of Cas9 RNP-loaded ARMMs across more than 10 different cancer, embryonic, and primary mouse and human cell types yielded productive editing with multiple gRNAs. Our data suggest that the efficiencies of delivery are commensurate with the properties of the gRNA, access to the target locus in a specific cell type and ARMMs uptake efficiency. Notably, all cell types showed appreciable editing, including primary post-mitotic cells.

Intranasal administration of ARMMs led to marked functional uptake in alveolar macrophages as well as Cas9-based editing of the Nlrp3 and Irf5 genes. Similarly, a high level of functional payload delivery to Kupffer cells, LSECs and splenic macrophages was attained through intravenous administration, indicating that ARMMs are potentially useful to target axes of inflammation or tolerance in macrophages. To illustrate the therapeutic potential of ARMMs in this context, we targeted the NLRP3 gene, which is highly expressed in Kupffer cells particularly in the disease state ^29^, and demonstrated in an ALI model significantly diminished pathology. While NLRP3 is a target that is “druggable” using small molecules, other well validated targets with therapeutic potential, such as the transcription factor IRF5, are not. ARMMs permit unconstrained targeting of any gene in Kupffer cells or alveolar macrophages using genome editing payloads, expanding the potential applicability of this platform to treat complex immune diseases, while limiting deleterious effects associated with broader inhibition of some targets. Furthermore, that ARMMs are potentially redosable increases the utility of this system in treating short-lived cells such as Kupffer cells.

Delivery of genome editors with ARMMs bestows several advantages over viral and other synthetic or virus-derived systems. First, as engineered forms of naturally existing vesicles in vertebrates, ARMMs are well tolerated even when human cell-derived vesicles are injected intravenously in mice. Second, because ARMMs deliver Cas9/gRNA RNPs which are prone to rapid intracellular turnover, it is probable that off-target editing, while not evaluated in this manuscript, is more limited than other systems. This is further ensured by the tethering of ARRDC1 to Cas9 which limits the number of payloads reaching the nuclear compartment ascertaining minimal editing only at the target locus defined by the gRNA of choice. Third, ARMMs are not subject to endosomal trapping due to the ability to overcome compartmentalization through the introduction of fusogens, which is not currently achievable in synthetic nanoparticle systems and often a source of dose-limiting toxicity ^30^. Last, ARMMs extend therapeutic potential to Cas9 RNPs in targeting genes expressed in macrophages, promoting evaluation of previously untested paradigms for a range of diseases. Evaluation of ARMMs in relevant models of disease is required to fully realize the therapeutic potential of these human cell-derived delivery vehicles.

## Materials and Methods

### Molecular cloning and plasmids

ARRDC1-Cas9 expression construct was made by cloning full-length DNA fragment of wild-type ARRDC1 and Cas9 with a G4S linker at the C-terminus of ARRDC1 into the pcDNA3.1(+) vector at Vectorbuilder. HEK3 gRNA plasmid was a gift from Dr. David Liu of Harvard University. ARRDC1-GFP and ARRDC1-CRE constructs were made in the pLV vector with CMV promoter and puromycin selection marker at Vectorbuilder, and the relative lentivirus was also made by Vectorbuilder. ARRDC1-mCherry construct was made in the JumpIn vector with puromycin selection marker at Vectorbuilder. All the other gRNA plasmids were made by Vectorbuilder. All gRNAs are expressed under the promoter of U6. New constructs were confirmed by direct DNA sequencing.

### Cell culture

HEK293T cells (ATCC; CRL-3216), NIH3T3 cells (ATCC; CRL-1658), Raw264.7 cells (ATCC; TIB-71) and sNF02.2 cells (ATCC; CRL-2885) were maintained in DMEM (ATCC; Catalog No. 30-2002) supplemented with 10% (v/v) heat-inactivated fetal bovine serum (FBS). Neuro-2a cells (ATCC; CCL-131) were maintained in EMEM (ATCC; Catalog No. 30-2003) supplemented with 10% (v/v) heat-inactivated FBS. A549 cells (ATCC; CCL-185) were maintained in F-12K Medium (ATCC; Catalog No. 30-2004) supplemented with 10% (v/v) heat-inactivated FBS. All cell lines were sub-cultured twice per week at the split ratio of 1:3 – 1:20, adjusted based on cell types and cell density. Cells were cultured at 37°C with 5% carbon dioxide and were confirmed to be negative for mycoplasma by testing with MycoStrip™ - Mycoplasma Detection Kit (InvivoGen; rep-mys-50) or Myco-Sniff Mycoplasma PCR Detection Kit (MP Biomedicals; 093050201).

Expi293 suspension culture was maintained in Expi293 Expression Medium (Thermo Fisher Scientific, A1435102) at 37°C with 5% carbon dioxide on a rotator at 250 rpm. Cells were sub-cultured twice per week with seeding density of 0.5-1×10^6^ cells/mL.

### ARMMs production, purification, and characterization

Expi293 stable cells expressing A1-GFP or A1-mCherry were seeded at a density of 0.5×10^6^ viable cells per mL and expanded for 4-5 days to a VCD of ∼5×10^6^/mL. Cells and large cellular debris were removed by centrifugation steps at 500 xG for 3 minutes and 2000 xG for 10 minutes, respectively. The clarified conditioned media (CM) were then 0.2 µm filtered and stored at 4°C until downstream processing.

Other ARMMs were produced by transiently transfecting wild type Expi293 cells using Expifectamine (Fisher Scientific, A14524) according to the manufacturer’s recommendations, omitting the Enhancer. Cells were transfected at a density of 3×10^6^/mL and allowed to recover overnight. The next day, cells were pelleted and re-suspended in fresh medium. Medium was conditioned for 3 days before clarification and storage as with the stable cells’ CM.

To purify the ARMMs, the CM was processed by tangential flow filtration (TFF) followed by ultracentrifugation. TFF was done on a Repligen KR2i system using a mPES hollow fiber filter with a 300 kD molecular weight cutoff, either microKros or midiKros format. The material was concentrated 5 times and washed with 5 continuous diavolumes of PBS, then further concentrated up to 15x. The retentate was 0.2 µm filtered, and then ultracentrifuged at 100,000 xG for 2h at 4°C in a Beckman Coulter Optima XE-90 ultracentrifuge using the swinging bucket rotor SW32 Ti. The supernatant was decanted and the tube briefly rested upside down to remove supernatant from the sides of the tube. Then the pellet was re-suspended in 200-500 µl PBS and sterile filtered twice using a syringe filter (Millipore Sigma’s Millex-GV PVDF 13 mm 0.2 µm filter, #slgv013sl). ARMMs were stored at 4°C.

To quantify the concentration and size of particles, NTA was done using the Particle Metrix Zetaview with the following settings: Sensitivity 82.4, shutter 100, minimum area 10, maximum area 3000, and minimum brightness 20. The particles were diluted in distilled water such that there were 50 – 200 on the screen, which is within the instrument’s linear range of detection.

### Western blot analysis and ARMMs payload quantification

1×10^9^ to 1×10^10^ ARMMs were lysed in Invitrogen Novex LDS Loading Buffer (Thermo Fisher Scientific, B0007) with β-mercaptoethanol included (5% v/v), and heated at 95°C for 10 minutes. The proteins were separated on a Bolt 4-12% acrylamide Bis-Tris gel (Thermo Fisher Scientific NW04125BOX) and transferred to a nitrocellulose membrane using the Trans-Blot Turbo system (Bio-Rad). The membrane was blocked with 5% milk in TBS-T for 1 hour at room temperature, and probed with primary antibodies to ARRDC1 (provided as a gift from Dr. Qian Lu, 1:5000), Cas9 (Cell Signaling Technology 14697S, 1:1000), syntenin (Abcam, ab133267, 1:1000), and vinculin (Cell Signaling Technology 13901S, 1:1000) overnight at 4°C with gentle rocking. The membrane was then washed 3 times for 5 minutes each in TBS-T and probed with a HRP-conjugated secondary antibody to mouse or rabbit IgG as appropriate (Cell Signaling Technology, 7076S or 7074S), at a dilution of 1:2000 for 1h with gentle rocking at room temperature. The membranes were then washed 4 times for 5 minutes in TBS-T, and developed using Pierce Western Blotting Substrate (Thermo Fisher Scientific, 32106) with imaging on an iBright imaging system (Thermo Fisher Scientific). When required, we stripped the membranes using Restore Western Blot stripping buffer (Thermo Fisher Scientific PI46430).

To calculate the cargo protein load in ARMMs, Western blotting was carried out with known amount of recombinant proteins (ARRDC1 or Cas9), from which a 6-point standard curve was generated. Concentrations for ARMMs samples were extrapolated from the standard curve and used to calculate copy number per ARMM using the particle concentration from NTA and the molecular weight of ARRDC1 or Cas9 to determine the number of molecules/particle.

### *In Vitro* ARMMs.mCherry.P Uptake Assay

Various types of cells were seeded at 25000 cells per well in 24-well plates and treated at the desirable concentrations of ARMMs on the same day. After 24 h, cells were trypsinized and washed with PBS three times before re-suspended in eBioscience Flow Cytometry Staining Buffer (Fisher Scientific, cat #50-112-9748) for flow cytometry analysis on Attune NxT (Thermo Scientific). For imaging, cells were seeded in 96-well plates at 10000 cells per well and treated with ARMMs at the desirable concentrations. After 24 h, live cells were imaged on Opera Phenix high content imaging system (PerkinElmer).

### HEK293 Cre reporter assay

HEK293-loxP-GFP-RFP cells (GenTarget, cat# SC018-Bsd) were plated at 15,000 cells per well in a 96-well plate in 150 µl. ARMMs were applied in a final volume of 50 µl to each well at the indicated concentrations, and cells were incubated for 24h. Cells were then washed once with PBS, released with 100 µl TrypLE for 5 minutes, and transferred to a V-bottom 96-well plate. The wells were washed with 100 µl of medium which was combined to wells in the V-bottom plate. The plate was centrifuged at 500 xG for 5 minutes, and the supernatant was pipetted off. The cell pellets were re-suspended in 200 µl of eBioscience Flow Cytometry Staining Buffer (Fisher Scientific, cat #50-112-9748) and analyzed by flow cytometry using Attune NxT (Thermo Scientific).

### ARMMs uptake and CRISPR/Cas9 genome editing evaluation

Cells were seeded in 96-well plates at the densities of 5000 cells/well for cell lines or 10,000 cells/well for mouse primary cells. After 3 hours, ARMMs were applied to cells at the indicated concentrations (ARMMs/cell). Cells treated for 96 hours were collected for genomic DNA (gDNA) extraction or stored at -80C for later processing. gDNA was extracted using the DNeasy 96 Blood & Tissue Kit (Qiagen, 69581) and following the manufacturer’s instructions. Targeted PCR was performed to amplify the genomic regions harboring the genome editing sites. 5 uL of gDNA solution was used as template in the 50 uL PCR reaction using Q5 High-Fidelity 2X Master Mix (NEB, M0492) with the manufacturer’s protocols. PCR products were confirmed on E-gels and purified using the QIAquick 96 PCR Purification Kit (Qiagen, 28181). 1-5 uL of the eluted PCR products were submitted for Sanger sequencing to have an initial evaluation of editing before the rest of PCR products were submitted for Amplicon deep sequencing.

### Quantitative PCR

Cells treated with ARMMs in 96-well plates were collected and stored at -80°C. RNA samples were prepared using TaqMan™ Gene Expression Cells-to-CT™ Kit (ThermoFisher Scientific, AM1728). Briefly, 50 uL of cell lysis buffer with DNase I (100X) was dispended to each well with pipette tips scraping well bottom to dislodge cells followed by incubation at room temperature (RT) for 5 min. 5 uL STOP solution from the kit was dispensed to each well to stop the lysis. The High-Capacity cDNA Reverse Transcription Kit (ThermoFisher Scientific, 43-749-66) was used for cDNA synthesis with 13 uL of the above-prepared RNA samples as template in a 20 uL reaction. At the end of the reaction, cDNA samples were diluted 4-fold. For qPCR reaction in 384-well plate, 10 uL reaction was assembled with 5 µL of TaqMan 2X Gene Expression Master Mix (Thermo Fisher Scientific, 4369016), 0.5 µL of 20X TaqMan primer probe for mouse Hprt gene (VIC), 0.5 µL of 20X TaqMan primer probe for the gene of interest (FAM), and 4 µL of prepared diluted cDNA. qPCR was run on QuantStudio Pro (ThermoFisher Scientific) and Cq values were used to evaluate gene expression levels normalized to the internal control *Hprt* expression.

### Evaluation of gRNA in ARMMs

To quantify gRNA packaged in ARMMs, a custom Taqman probe was ordered from ThermoFisher Scientific (Assay ID: AP7DWZ9). This probe was designed against the scaffold sequence in the single gRNA (sgRNA). The quantification of gRNA included three steps: preparation of RNA samples, cDNA synthesis, and droplet digital PCR (ddPCR). RNA samples were prepared using TaqMan™ Gene Expression Cells-to-CT™ Kit (ThermoFisher Scientific, AM1728). 1 uL of ARMMs preparation was added to 21.5 uL of cell lysis buffer with DNase I (100X), of which the mixture was incubated at RT for 5 min before 2.5 uL of STOP solution was added, resulting in 25X dilution of the original ARMMs preparation. Reverse transcription (RT) was performed with the 20X RT enzyme mix and 2X RT buffer from the kit in a 25 uL reaction with 1 uL of prepared RNA samples. Droplet digital PCR was assembled in a 25 uL reaction as follows: ddH2O, 17 uL; 5X Naica multiplex PCR mix, 5 uL; 100% Buffer B, 1 uL; gRNA scaffold Taqman probe (25X), 1 uL; cDNA, 1 uL. Assembled ddPCR mixture was loaded to Naica chips and run on the Naica™ System for Crystal Digital PCR™ (Stilla Technologies). The relative amount of gRNA in ARMMs was reported as fold enrichment by normalization to the control ARMMs produced with Cas9 plus HEK3 gRNA without ARRDC1 fusion.

### Blood and tissue processing and mCherry quantification by ELISA

Whole blood was collected from each mouse in tubes coated with EDTA to avoid clotting. The blood was centrifuged at speed of 800g for 5min at room temperature, and plasma was transferred to a new Eppendorf tube for mCherry quantification. All major organs were harvested from each mouse and flash frozen using liquid nitrogen. The tissues were then pulverized using the 1600MiniG Spex SamplePrep. 1x Cell Extraction Buffer PTR (Abcam, Cambridge, MA) was added to the pulverized tissues and homogenized again. Protein isolation was done by centrifuging the samples at 15000g for 20min at 4 degree C and protein concentration was measured in the supernatant using Pierce™ Rapid Gold BCA Protein Assay Kit (Thermo Fisher Scientific, 23227). Protein quantifications were performed following manufacturer instructions and biological samples were prepared under the same conditions as standards. Extracted protein concentrations were calculated and all samples were normalized to 6mg/ml. All plasma samples were diluted in the 1x Cell Extraction Buffer PTR for the assay. mCherry protein levels were then quantified using an mCherry-specific ELISA kit (Abcam, ab221829).

### Mouse strains and housing

Female C57BL/6 mice (strain code #027, 6-8 week) were purchased from Charles River Laboratories, and female Ai9 mice or Ai14 mice were purchased from Jackson Labs (strain #007909 or #007914, respectively). In the case of acute liver injury study, male C57BL/6 mice were used (Charles River Laboratories, strain code #027, 6-8 week). Mice were housed in Charles River Vivarium (Cambridge, MA) and standard mouse husbandry procedures were followed. All animal experiments were approved by the Animal Care and Use Committee of Charles River Laboratories.

### Mouse primary cell Isolation

To isolate liver cells including hepatocytes, Kupffer cells and liver sinusoidal endothelial cells (LSEC), paranchymal and non-paranchymal liver cells were isolated by two-step liver perfusion. Briefly, livers were perfused through the inferior vena cava with 1mM EDTA, 25mM HEPES in PBS followed by perfusion with 1mg/mL Collagenase IV (Sigma C428-100MG) in DMEM. Livers were then excised from the mouse and nicked to release cells. Cell suspension was then filtered through a 70um cell strainer. Hepatocytes were isolated by gentle centrifugation at 40 xg for 5 min at 4°C and plated at 2×10^4^ cells per well in 96-well tissue culture plates coated with 0.1mg/mL rat-tail collagen (Corning 354236) and cultured in DMEM (Gibco 10564-011) with 5% FBS (Gibco 10082147) and 1xPen/Strep (Gibco 15070063). Non-paranchymal cells in the supernatant were then spun at 300 xg for 10min at 4°C. Kupffer cells were isolated using F4/80 microbeads (Miltenyi Bio 130-110-443) according to manufacturer’s instructions and cultured in DMEM with 5% FBS and 1xPen/Strep. Liver sinusoidal endothelial cells were isolated using CD146 microbeads (Miltenyi Bio 130-092-007) according to manufacturer’s instructions and cultured in EBM-2 media with supplements (Lonza CC-3162). LSECs were plated at 2×10^4^ cells per well in 96-well tissue culture plates. Primary hepatocytes were used in *in vitro* gene editing experiments. LSEC were used in *ex vivo* Cre uptake experiments.

Lung alveolar macrophages were isolated with different methods for different experiments. For ex vivo Cre uptake experiments or flow cytometry analysis of *in vivo* ARMMs uptake experiments, alveolar macrophages were isolated by lung digestion as previously described ^31^. Lungs were inflated *in situ* with 1mL of digest media #1 (RPMI (Gibco 12633-012) containing 4 U/mL elastase (Worthington #LS002290), 1 U/mL Dispase (Sigma #D4693), and 200 ug/mL DNAse (Sigma #10104159001)) using a gavage adaptor for a 1-mL syringe and trachea clamped to prevent leaking. The lung was excised from the mouse and placed into 2-mL of digest media #1 for 30min in a 37°C water bath. The lung was then removed from the digest media and cut into small pieces followed by a second digestion step in digest media #2 (5mL RMPI, 25ug/mL liberase (Sigma 05401020001) and 200ug/mL DNAse) for 30 min in a 37°C incubator. The cell suspension was then filtered through a 70um cell strainer and pelleted at 400 xG for 5 min at 4°C followed by RBC lysis using ACK lysis buffer (Gibco A10492-01) for 5 min on ice. Single cell suspensions were then used for downstream flow cytometry analysis or lung macrophages were isolated from single cell suspension by selection with CD11c microbeads (Miltenyi 130-125-835) according to manufacturer’s instructions for *in vitro* analysis. Macrophages were plated at 2×10^4^ cells in 96-well tissue culture plates and cultured in DMEM with 10% FBS. For *in vivo* gene editing experiments, alveolar macrophages were isolated from bronchoalveolar lavage fluid (BAL). Lungs were washed 10 times with 1-mL of buffer (PBS containing 1% FBS and 2mM EDTA) using a gavage adapter to perfuse buffer down the trachea. BAL was centrifuged at 400 x g, followed by RBC lysis. Cell pellets were frozen and used for downstream gene-editing analysis.

To isolate skin fibroblasts, shaved skin free of fur was excised from the underarm area of mice with caution to not isolate the fat layer along with the skin. Skin was then cut into 1mm pieces using two scalpels to resemble a putty-like consistency. Skin fragments were then incubated in 4mL of 2mg/mL Collagenase Type I (Sigma SCR103) in DMEM (Gibco 10564-011) in a 37°C water bath pipetting up and down each 5 min during the incubation to break apart clumps. After incubation, cells and tissue fragments were pelleted at 500 xG at 4°C for 5min and resuspended in 10mL of fresh plating media containing DMEM (Gibco 10564-011), 1xPen/Strep (Gibco 15070063), and 5% FBS (Thermo 10082147). Cells and tissue fragments were plated in a 10cm dish and incubated at 37°C, 5% CO2, 3% O2 for 5 after which the plate was washed three times with plating media. After reaching confluence, skin fibroblast were plated at 1×10^4^ cells/well in a 96-well plate in plating media overnight. The media was then changed to serum-free plating media to prevent proliferation of skin fibroblast for 24 h before addition of ARMMs.

To isolate bone marrow derived macrophages (BMDM), freshly collected bone marrow cells were flushed from the femur and tibias with PBS, filtered through 40um cell strainer, and cultured in DMEM (GIBCO Cat#10-564-011) containing 10% HI FBS (GIBCO Cat#10082147) and 50ng/ml M-CSF (R&D Systems Cat# 416-ML-010) for 7 days and re-plated at 20,000 cells per well in 100 µl DMEM containing 10% FBS in 96-well microplates. Cells were allowed to recover overnight before ARMMs treatment.

### Staining and quantification of functional delivery of ARMMs.Cre.P to primary cell cultures

ARMMs.Cre.P were added to primary cell cultures of liver sinusoidal endothelial cells, lung macrophages, skin fibroblasts, and bone marrow derived macrophages at concentrations of 1×10^4^, 1×10^5^, and 1×10^6^ ARMMs/cell directly to cell culture media in each well. As a negative control, PBS was added to primary cells. Cells were incubated with ARMMs for 72-h before fixation and quantification of uptake. Three replicates per concentration were performed for each cell-type analyzed.

Wells were washed twice with Phosphate buffered saline and then fixed with 4% Paraformaldehyde for 1 h at room temperature. Cells were permeabilized with 0.4% Triton X (Sigma-Aldrich T9284) in PBS for 15 min at room temperature. Wells were briefly washed in PBS followed by 1 h incubation with animal free blocking media (Vector Labs SP-5035). Following blocking, cells were incubated with primary antibody diluted in PBS overnight at 4°C at the following concentrations: Rabbit anti CD206 (Cell Signaling 24595S, 1:200), Rabbit anti LYVE-1 (Abcam ab218535, 1:1000), Rabbit anti Smooth muscle actin (Abcam ab32575, 1:400). Wells were washed three times with PBS and incubated with secondary antibody (Abcam ab150077 Goat anti Rabbit 488, 1:500) for 2 h at room temperature. Following incubation, wells were washed three times with PBS followed by incubation with DAPI nuclear stain (1:2000 dilution of Abcam ab228549) for 10min at room temperature. Wells were then imaged for DAPI, Cell-markers, and tdTomato using an ImageXpress automated imaging instrument.

tdTomato positive cells from primary cell culture experiments were quantified in ImageJ. At least four fields of view from three replicates (n > 600 cells) were quantified for each condition. DAPI stain was used to determine total number of cells per field of view. To determine tdTomato positive cell number, images were thresholded to display tdTomato positive signal and cell number counted. The ratio of tdTomato positive cells to total nuclei number per field of view was used to calculate percentage of tdTomato positive cells.

### Mouse dosing

Mice were dosed by oropharyngeal aspiration with 50uL of solution containing 5×10^11^ particles or 50uL of Phosphate Buffered Saline (PBS). Briefly, mice were anesthetized via isoflurane, and front teeth (incisors) placed over a silk suture suspended across an angled platform to facilitate access to the oral cavity. The tongue was pulled from the oral cavity using blunt forceps to prevent the swallowing reflex and the nose of the mouse was covered to facilitate inhalation through the mouth. 50uL of test article solution was then pipetted into the mouse’s mouth. The mouse was held in the described position until an audible inhalation through the mouth was heard. The mouse was returned to its cage and monitored until waking from anesthesia.

Mice dosed intravenously with ARMMs were given a single retro-orbital injection containing 1×10^12^ particles or 200uL of Phosphate Buffered Saline (PBS) after being anesthetized via isoflurane.

For ARMMs payloaded with Cre or Cas9-gRNA RNP, lungs or livers were collected from mice 96 h after administration for downstream processing. For ARMMs payloaded with mCherry, lungs were collected from mice 2 h after administration for downstream processing, or at prolonged time points when the time course of ARMMs.mCherry.P uptake was examined.

### Flow cytometry analysis of *in vivo* alveolar macrophage, Kupffer Cell, and Liver sinusoidal endothelial cell ARMMs uptake

Approximately 5×10^5^ to 1×10^6^ cells isolated from lung or liver digestion were used for flow cytometry staining. Prior to antibody staining, live/dead fixable near IR (Thermofisher L10119) staining was carried out according to manufacturer’s instructions.

Alveolar macrophages from lung digestion were identified as CD45^+^ CD11b^int^ CD11C^hi^ SiglecF^+^. The following antibodies were used at a 1:100 dilution in Flow Cytometry Staining Buffer (eBiosciences #00-4222-26): CD45-PerCP Cy5.5 (Biolegend 103132), CD11b-Alexafluor 488 (thermofisher 53-0112-82), CD11c-APC (Biolegend 117310), and SiglecF-PE/Cy7 (Biolegend 155527).

Liver sinusoidal endothelial cells were identified as CD45^-^CD31^+^CD146^+^ and liver Kupffer cells were identified as CD45^+^CD11b^mid^F4/80^+^. The following antibodies were used at a 1:100 dilution in Flow Cytometry Staining Buffer (eBiosciences #00-4222-26): CD45-PerCP Cy5.5 (Biolegend 103132), F4/80-FITC (Biolegend 157310), CD11b-APC (Biolegend 101212), CD31-Alexafluor700 (Biolegend 102444), CD146-PE-Cy7 (Biolegend 134714)

mCherry or tdTomato intensity in the alveolar macrophage population was evaluated by native fluorescence. mCherry or tdTomato positive cells were gated based on mice administered PBS in place of ARMMs as a negative control. Flow cytometry was performed on an Attune NxT flow cytometer, and data analysis performed on FCS Express (version 7).

### Immunohistochemistry of tissue sections

Lungs were isolated and each lobe separated individually and fixed in 4% PFA overnight at 4°C. For livers, either the left lateral lobe or the caudate lobe was isolated and fixed in 4% PFA overnight at 4°C. For paraffin processing, samples were transferred to 70% ethanol and processed by Alamak Biosciences (Beverly, MA) and sectioned at 5um thickness. For Frozen section processing, samples were transferred from 4% PFA to 15% sucrose for 8 h, followed by a transfer to 30% sucrose overnight at 4°C. Tissues were then embedded in Optimal Cutting Temperature media (Tissue-Tek O.C.T) and sectioned on a Leica CM1860 cryostat at 10um thickness and stored at -80°C.

For histology, tissue cross sections of Lung left lobes and liver caudate lobes were assessed for payload delivery. For immunofluorescence staining of paraffin embedded tissues, sections were deparaffinized in Xylene for 2 washes of 10 min, followed by a descending series of alcohol washes of 100%, 95%, 70%, and 50% ethanol followed by two washes in water. Antigen retrieval was then performed using Vector Labs citrate-based antigen unmasking solution (Vector Labs H3300) according to manufacturer’s instructions. Frozen sections were dried at room temperature for 20 min and washed in PBS for an additional 10 mins. Both paraffin and frozen sections were then permeabilized in 0.4% Triton-X in Phosphate-buffered saline. Sections were briefly washed in PBS followed by 1 h incubation with animal free blocking media (Vector Labs SP-5035). Sections were once again washed in PBS before incubation with primary antibody overnight at 4C.

For lung cross section immunostaining, primary antibodies used were rabbit-CD206 (Cell Signaling 24595S) and chicken-mCherry (Aves MCHERRY-0100) at 1:250 dilution in PBS. For liver lobe immunostaining, primary antibodies used were rabbit-Lyve1 (Abcam ab218535) at a 1:2000 dilution in PBS or rabbit-IBA1 (abcam ab178847) used at a 1:400 dilution in PBS. Sections were washed three times with PBS for 10 min each, followed by incubation with secondary antibody for 2 h at room temperature. Secondary antibodies used were Goat anti Chicken AlexaFluor 488 (Abcam ab15069) and Goat anti rabbit Alexafluor 647 (Abcam ab150075). For imaging of tdTomato signal in tissue cross sections, native fluorescence was observed from O.C.T embedded tissue cross sections.

Samples were washed three times for 10 min each in PBS before mounting with Vectashield Plus antifade mounting media with DAPI (Vectorlabs #H-2000). Images were taken on an EXC-500 microscope (Accu-scope) fitted with a DHYANA CMOS camera for fluorescent imaging (TUCSEN) and recorded using Mosaic imaging software (Tucsen, version 1.6).

### Acute liver injury (ALI) model

A single administration of 1×10^12^ particles (200 µL of 5 ×10^12^ particles/mL) of ARMMs.Cas9-Nlrp3.RNP or PBS vehicle control were delivered via tail vein intravenous injection to a group of 7 C57BL/6 male mice (6-8wk) to assess their effects three days prior to IV injection of 0.5 µL/g body weight CCl4 (Sigma-Aldrich, 319961-500ML diluted in corn oil [Sigma-Aldrich, C8267-500ML] at 2 µL/g) to elicit liver injury. Mice were euthanized one day after CCl4 treatment to assess their inflammation. Aliquots of 20 µL serum from terminal bleeds of mice were analyzed for liver damage markers AST and ALT (IDEXX). Hemolysis and lipemia index of these terminal bleeds were also assessed. Liver sections were collected for creation of single cell suspensions to assess in flow cytometry and tissue sectioning for immunofluorescence microscopy and H&E staining.

### Histology from ALI study

Histological slides (scanned, H&E-stained) containing liver were examined. The pathologist was blinded by the experimental groups (including controls), test-article structure, class, organ weights, and gross findings. Electronic slides were evaluated, and images were obtained, using the SlideViewer software. International Harmonization of Nomenclature and Diagnostic (INHAND) Criteria standards were used as the basis of histopathological evaluation (Thoolen et al 2010). The histopathology severity grades for hepatic necrosis are based on a previously described grading scheme (Shackelford et al, 2002) using the following criteria: 0= lesions are not present; 1= Minimal (when lesions involved < 1%); 2= Slight (when lesions involved 1–25% of the tissue section); 3= Moderate (when lesions involved 26–50% of the tissue section); 4= Moderately severe/high (when lesions involved 51–75% of the tissue section); 5=Severe/high (when lesions involved 76–100% of the tissue section). For necrosis-associated inflammation, the grading was based on the average number of leukocytes present in 10 x HPF (1 HPF=0.1123 mm2) from the areas of necrosis, using the following criteria: 0= if there are on average < 1 leucocyte / HPF; 1= Minimal, if there are on average 1-5 leucocytes / HPF; 2= Slight, if there are on average 6-10 leucocytes / HPF; 3= Moderate, if there are on average 11-20 leucocytes / HPF; 4= Moderately severe, if there are on average 21-30 leucocytes / HPF; 5= Severe, if there are on average > 31 leukocytes / HPF.

For quantification of apoptotic cells via TUNEL assay, paraffin embedded cross sections from the left lateral lobe were processed using one-step TUNEL apoptosis kit (Elabscience E-CK-A322) according to manufacturer’s instructions. Images from five regions (upper left, upper right, lower left, lower right, and center) were taken from two sections per mouse on an EXC-500 microscope (Accu-scope) fitted with a DHYANA CMOS camera for fluorescent imaging (TUCSEN) and recorded using Mosaic imaging software (Tucsen, version 1.6). TUNEL-stained images were thresholded in ImageJ software and the average percent thresholded area from the five regions was recorded as percentage TUNEL positive area. The average percentage of TUNEL positive areas from all regions and sections were calculated and reported for each individual mouse.

### TLR activation assay

THP1 cells were cultured in RPMI medium supplemented with 10% Fetal Bovine Serum with 5uM phorbol 12-myristate 13-acetate (Sigma-Aldrich P8139) for 3 days to differentiate them into macrophages. Cells were seeded at 20,000 cells/well in a 96 well tissue culture plate and treated with various doses of ARMMs and TLR agonists for 6 hours. The supernatant was then collected for LEGENDplex™ Human Inflammation Panel 1 (13-plex) (Biolegend 740808) analysis as per the manufacturer’s instructions.

Freshly isolated C57/B6 mouse bone marrow cells were differentiated into macrophages with 50ng/ml mouse recombinant m-CSF for 7 days. Macrophages were treated with TLR agonists or ARMMs for 6h. Supernatant from cell culture was analyzed for cytokine release.

### Neutralizing antibody study

To measure if ARMMs induce immune response, whole blood was collected from mice injected with ARMMs and the presence of neutralizing antibodies was evaluated by the blockage of ARMMs uptake in cultured A549 cells. Specifically, 10-week-old C57/B6 female mice were retrobulbar IV injected with ARMMs.GFP.P either with 1 dose or 3 doses of 1×10^12^ particles weekly with corresponding collection of whole blood in 1 week, 2 weeks or 3 weeks. Plasma was collected by centrifuged at speed of 800xg for 5min at room temperature. Plasma was stored at -80 C for further study. Neutralizing antibody competition assay was conducted as follows. A549 cells were plated at a seeding density of 20,000 cells per well in a 96 well tissue culture treated plate and cultured overnight in F-12K Medium (ATCC 30-2004). Collected C57/B6 plasma were first diluted 20-fold in media as working solution, and then followed by 2-fold serial dilutions. Media from each well was then replaced with 100ul of the prepared plasma titrations spiked with 1×10^5^ ARMMs.GFP.P. Plate was incubated at 37C for 4 hours. All cell wells were washed carefully with medium three times and then lysed with 1x Cell Extraction Buffer PTR. GFP protein levels were then quantified using a GFP ELISA Kit (Abcam, ab171581).

### Computational biology

#### Data analysis

To evaluate editing efficiency using Sanger sequencing data, Sanger data files along with the sgRNA target sequences were input into the command line Inference of CRISPR Edits (ICE) algorithm ^32^ to determine indel efficiency. To evaluate editing efficiency using Amplicon NGS sequencing, the fastq files were analyzed using batch mode in the Crispresso2 command line software ^33^ to determine indel efficiency with previously described parameters ^34^.

#### Design of gRNAs

CRISPOR command line software ^35^ was used to identify gRNAs and rank them according to in silico off-target and predicted efficiency scores (i.e., MIT specificity score^36^, CFD specificity score^37^, Doench ’16 score ^38^ and Moreno-Mateos score ^39^).

## Acknowledgements

We thank the graphic designers of Biopharma Dealmakers (Nature publishing group) for assistance with figure generation.

## Author contributions

J.F.N. conceived the project. Q.W., S.-L.L., W.-N.Z., R.V.K. and M.T. designed experiments. W.-N.Z., Z.B. and J.B. performed experiments related to *in vitro* editing, and *in vitro* ARMMs.mCherry uptake. S.-L.L., R.V.K. and C.O.S. performed *in vivo* and *ex vivo* experiments. Q.W., K.V. and C-W.C performed experiments related to NLRP3 and IRF5. Y.S. and P.M. performed experiments related to immunogenicity and biodistrution evaluation by ELISA. K.L., L.G., A.F., A.B. and S.G. contributed to ARMMs production. N.V. contributed to *in vitro* human cardiomyocyte experiments. C.C. contributed to computational data analysis. J.F.N. and W.-N.Z. drafted the manuscript with input from other authors.

---

**Supplementary Fig.S1.**
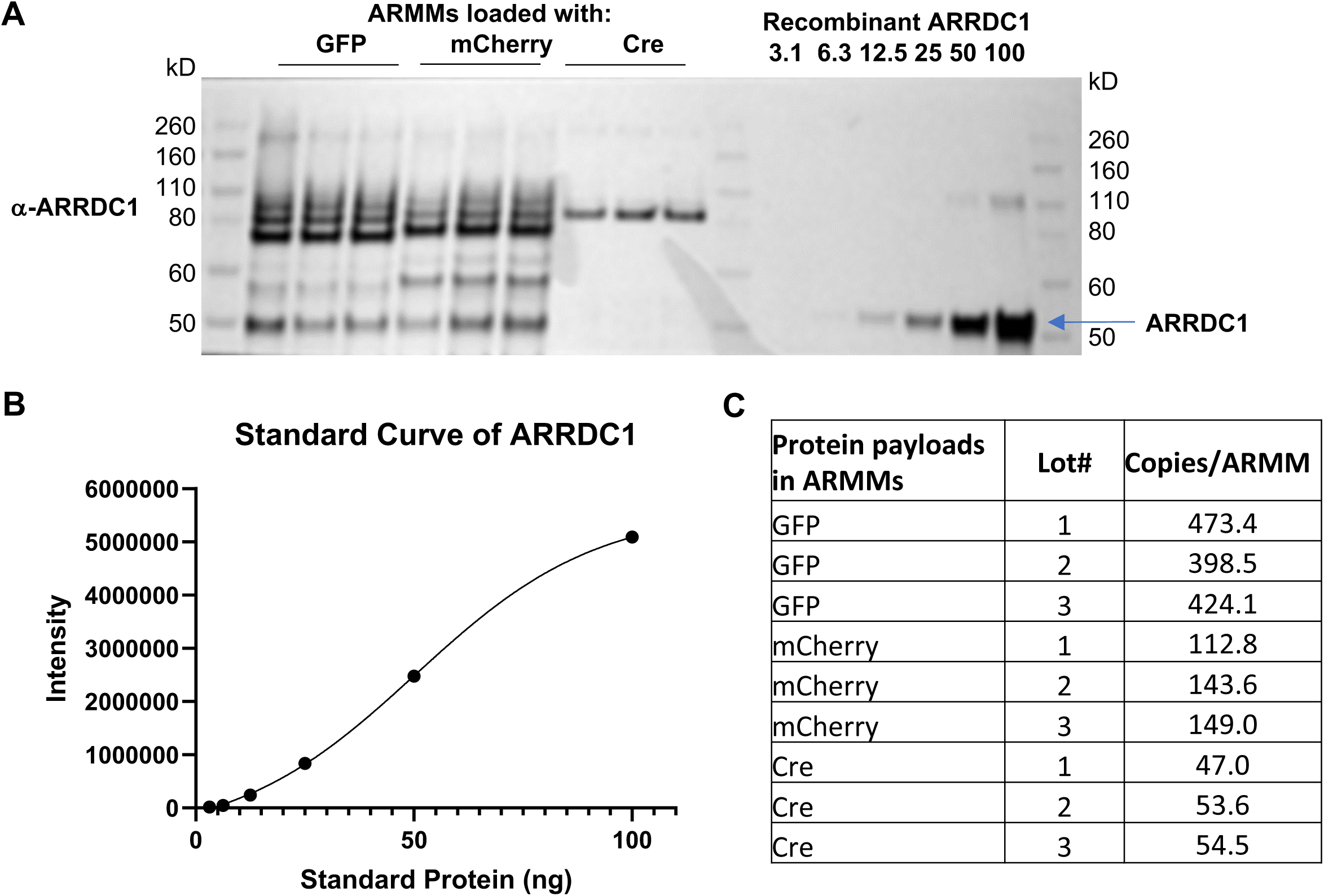
Evaluation and quantification of protein cargo in ARMMs. (**A**) Anti-ARRDC1 Western blot analysis of 3 batches of ARMMs (5×10^9^/well) loaded with GFP, mCherry or Cre proteins, and a serial dilution of human ARRDC1 recombinant protein. (**B**) A standard curve was generated from the recombinant ARRDC1 protein immunoblot. (**C**) Calculated copy numbers of GFP, mCherry or Cre per ARMM.

**Supplementary Fig.S2.**
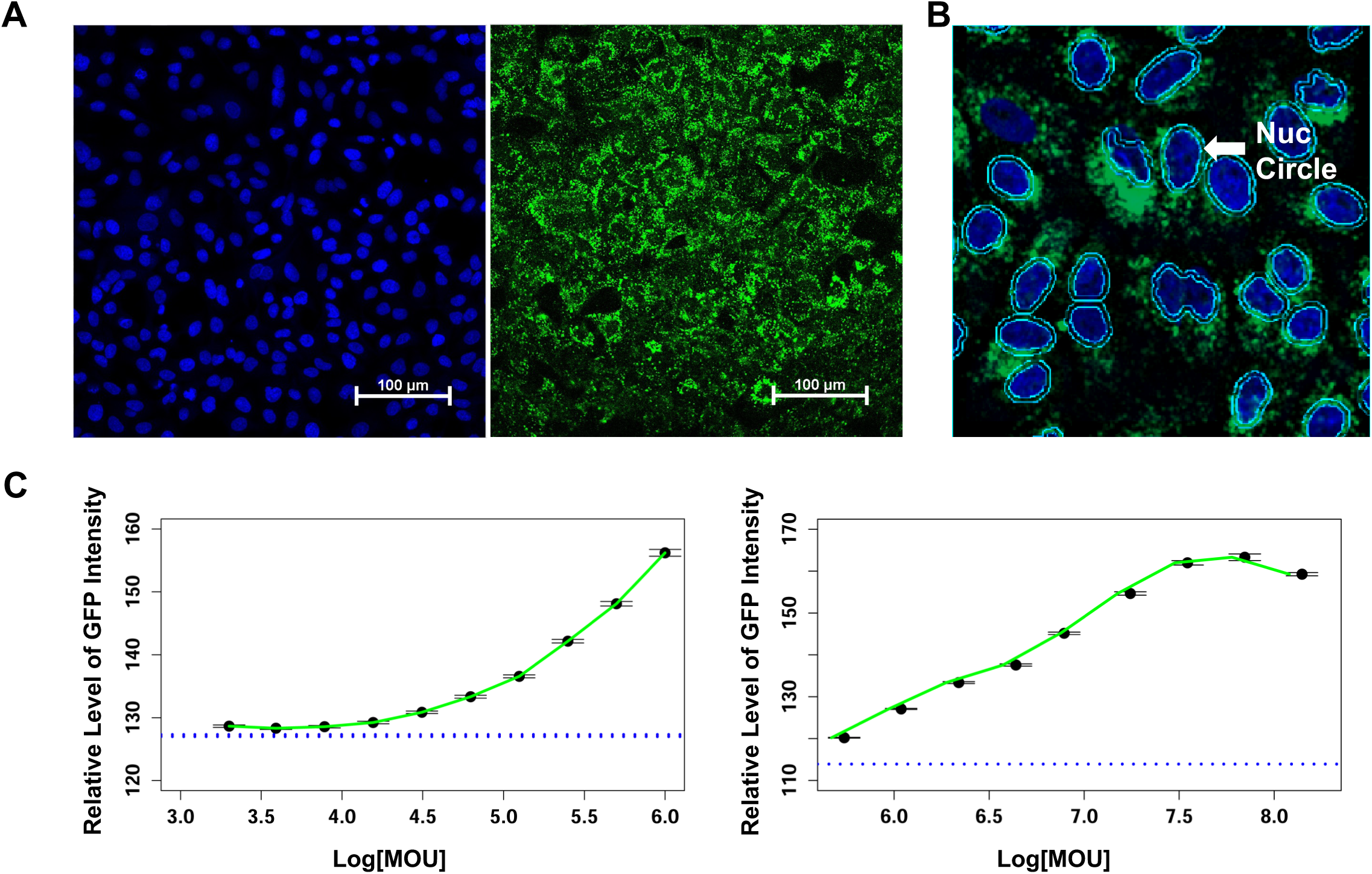
GFP protein delivery by ARMMs. ARMMs loaded with GFP protein (ARMMs.GFP.P) were applied to A549 cells and their uptake was recorded by fluorescence microscopy. (**A**) High content images from PBS or ARMMs.GFP.P treated A549 cells. DAPI was used to stain nuclei. (**B**) High content image analysis was used to quantify intracellular GFP protein delivered using ARMMs. Nuc circle defines the boundaries of the nucleus and extranuclear GFP signal was quantified. (**C**) Quantification of GFP signal after 24h treatment. The treatment concentrations ranged from 5×10^3^/cell to 1×10^8^/cell. ARMMs.GFP.P uptake was detected at a low MOU (multiplicity of uptake) of 1×10^4^ particles/cell and increased up to 5×10^7^ particles/cell.

**Supplementary Fig.S3.**
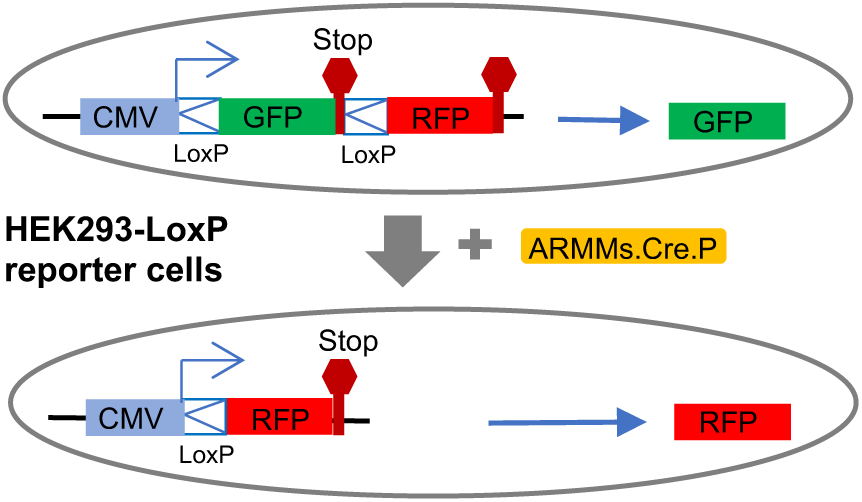
Illustration of Cre reporter cell line. GFP expression cassette is flanked by LoxP sites followed by an RFP expression cassette. In the absence of Cre recombinase activity, GFP is expressed, whereas in the presence of Cre recombinase, recombination happens at the two LoxP sites, leading to the excision of GFP expression cassette. As a result, RFP is expressed. The transition from GFP to RFP in cells is indicative of functional Cre activity.

**Supplementary Fig.S4.**
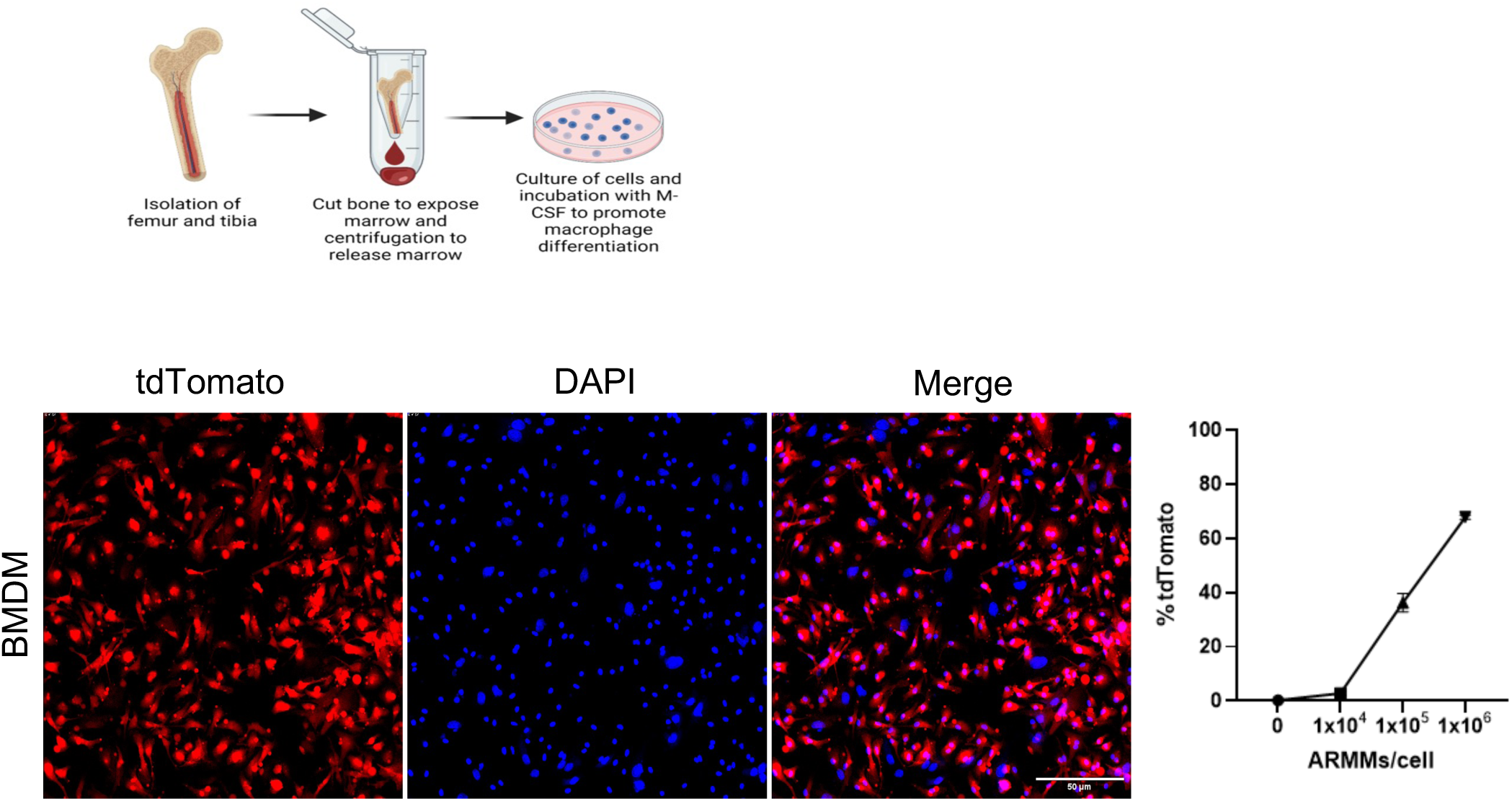
Functional uptake of ARMMs.Cre.P in mouse bone marrow derived macrophages (BMDM). (**A**) Depiction of methodology used for isolation of BMDM from Ai14 Cre-reporter mice. Bone marrow cells were harvested from hip bones and the sternum, and cultured in RPMI growth medium supplemented with 10% FBS and 50ng/ml recombinant mM-CSF for 3 days. The bone marrow-derived macrophages were plated at 20,000 cells/well in a 96 well plate and treated with a dose titration of ARMMs.Cre.P for 2 days. Cells were then washed with PBS three times and fixed with 4% paraformaldehyde (PFA) and nuclei were stained with DAPI. (**B**) Representative images showing tdTomato+ cells (left) and quantification of tdTomato+ cells upon treatment with increasing dose of ARMMs (right). Images were taken at 200x using ImageXpress nano and image analysis was carried out using MetaXpress software.

**Supplementary Fig.S5.**
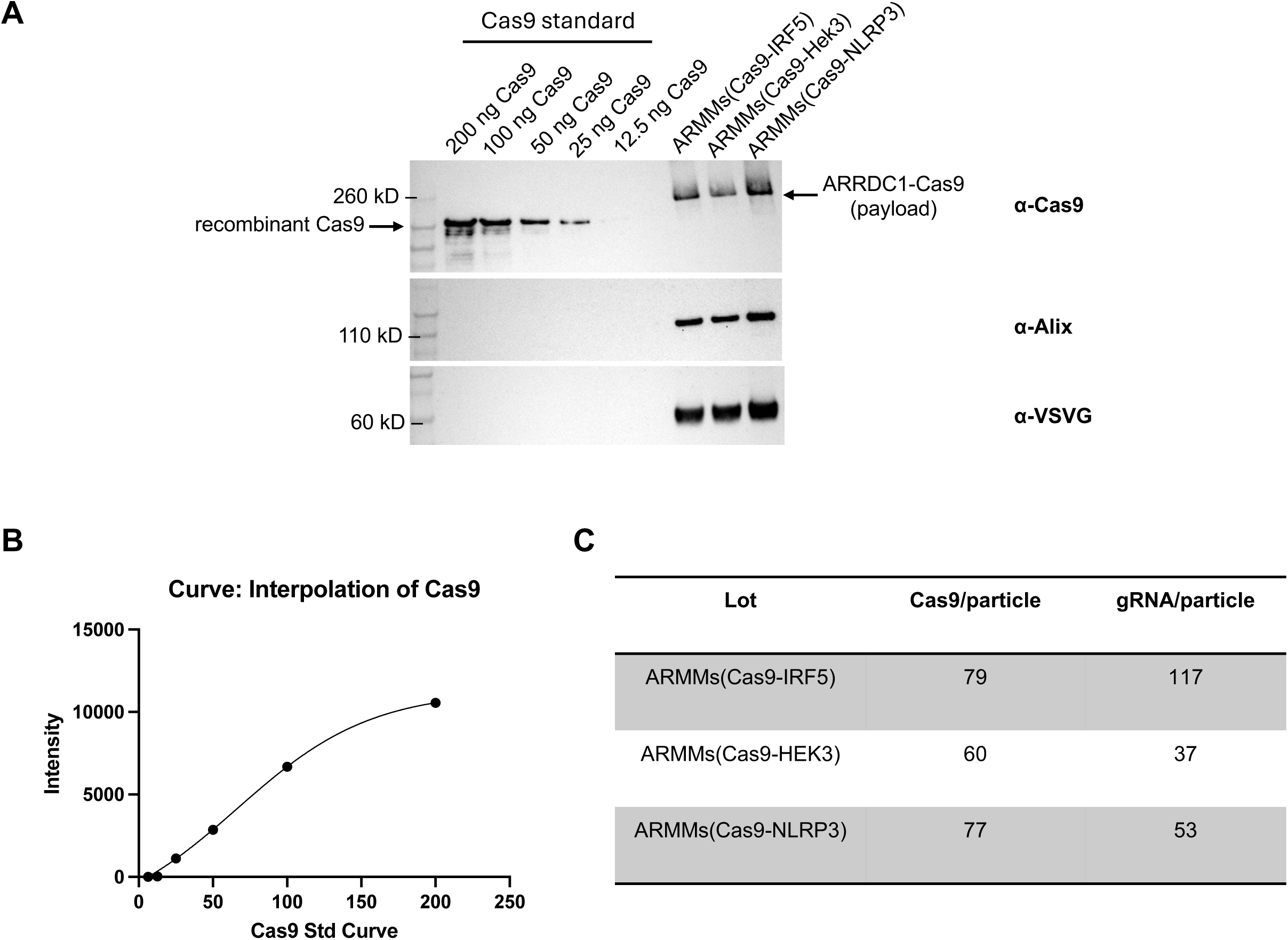
Evaluation of Cas9-gRNA payloads in ARMMs. (**A**) Western blotting to evaluate loading of Cas9 in ARMMs. 5×10^9^ ARMMs loaded with Cas9/gRNA RNPs from 3 batches were analyzed along with a serial dilution of recombinant Cas9 protein by immunoblotting with anti-Cas9 antibody. (**B**) Standard curve generated from densitometry analysis of the immunopositive bands corresponding to recombinant Cas9 protein standard. (**C**) Calculated copy numbers of Cas9 and gRNA per particle for purified ARMMs shown in A. gRNA evaluation was done by dPCR with a Taqman probe complementary to a common scaffold sequence in gRNAs.

**Supplementary Fig.S6.**
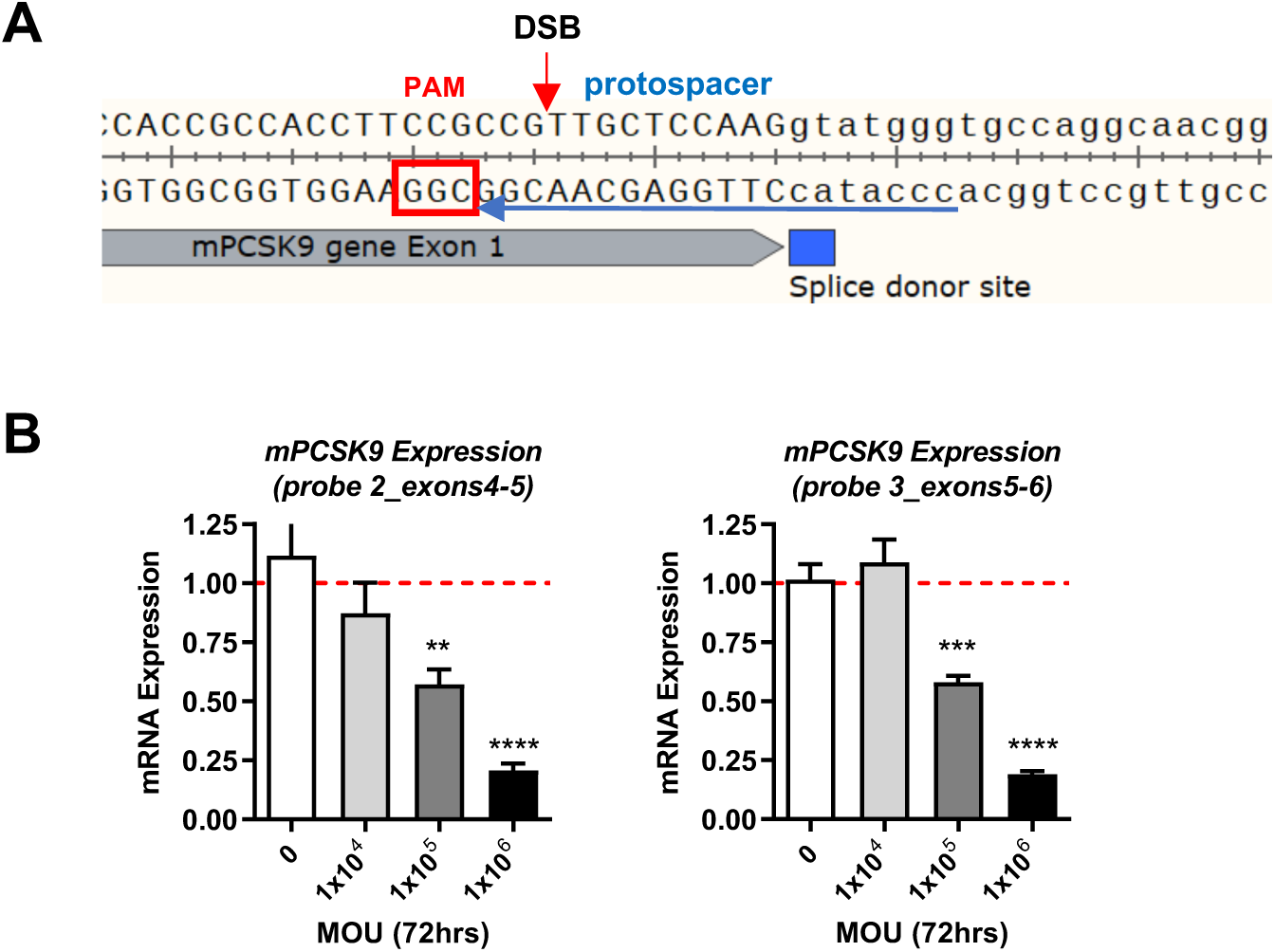
Treatment of mouse primary hepatocytes with ARMMs.Cas9-mPcsk9.RNP. (**A**) Illustration of sequence derived from Exon 1 of mouse Pcsk9 gene highlighting the gRNA target region/protospacer (blue underline), PAM (red box), and location of Cas9-induced double-stranded break (DSB). (**B**) Quantitive PCR data showing reduced expression of Pcsk9 mRNA upon treatment with ARMMs.Cas9-mPcsk9.RNP. Shown here are results from two additional Taqman probes targeting different regions of Pcsk9 mRNA to that shown in Fig.3F.

**Supplementary Fig.S7.**
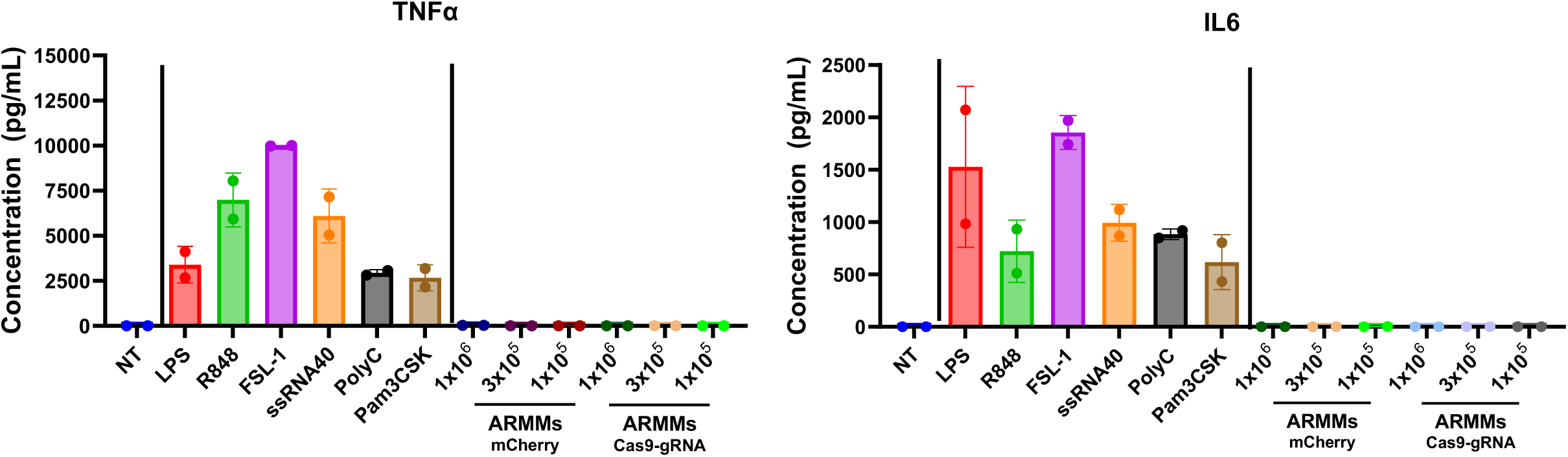
ARMMs do not activate Toll-like receptors (TLRs) in mouse bone marrow derived macrophages. Freshly isolated C57/B6 mouse bone marrow cells were differentiated into macrophages with 50ng/mL mouse recombinant m-CSF for 7 days. Macrophages were treated with TLR agonists and ARMMs for 6h. Supernatants from cell culture were analyzed for cytokine release. NT: not treated; LPS: 10ng/mL; R848: 10ug/mL; FSL-1: 25ug/mL; ssRNA40: 25ug/mL; PolyC: 10ug/mL; Pam3CSK: 25ng/mL.

**Supplementary Fig.S8.**
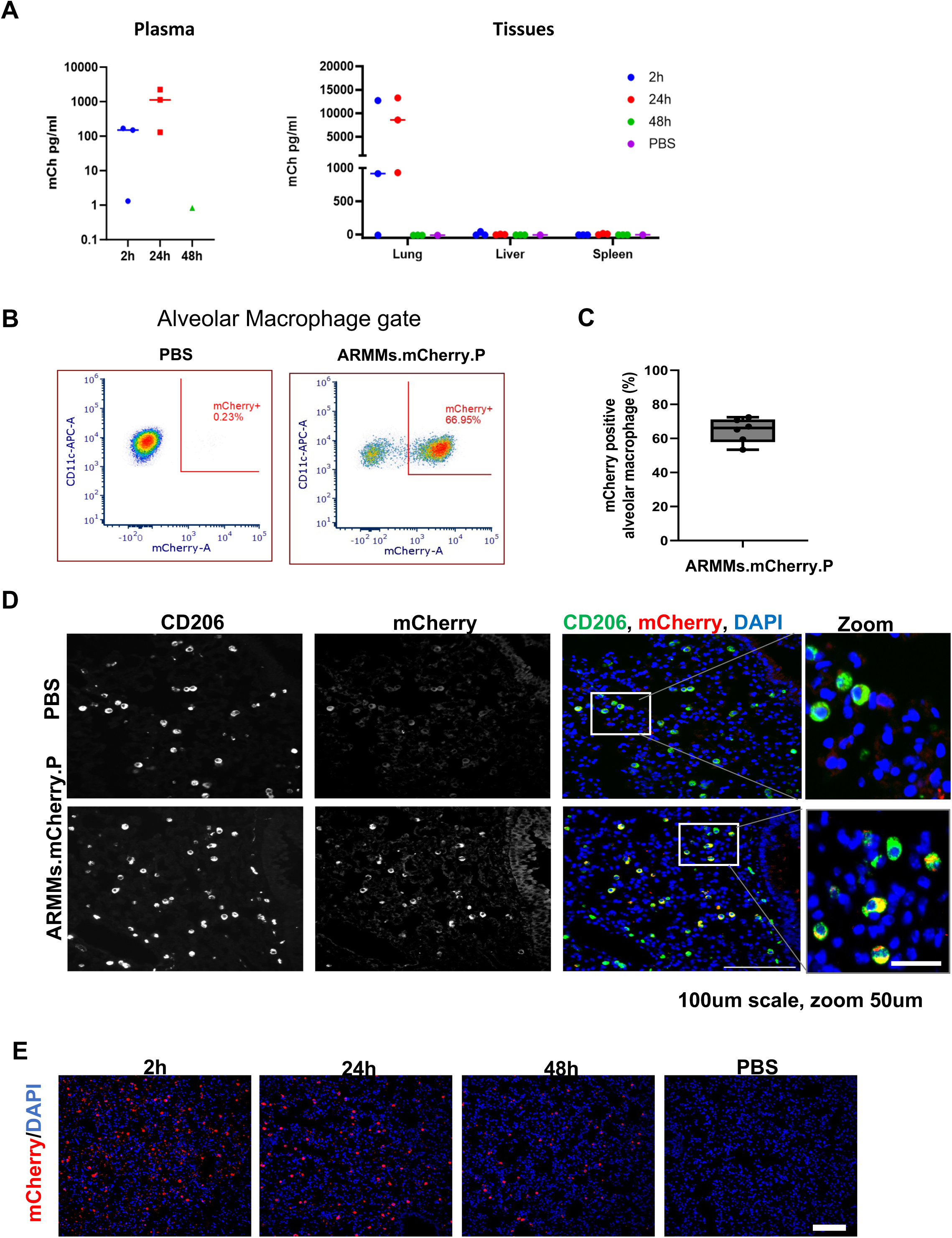
Oropharyngeal aspiration of ARMMs loaded with mCherry results in uptake in alveolar macrophages. **(A)** Pharmacokinetic evaluation of ARMMs in plasma, spleen, and liver. **(B)** Representative flow cytometry density plots of ARMMs.mCherry.P uptake in alveolar macrophages in a PBS or ARMMs-treated animal. mCherry-positive alveolar macrophages are boxed inside the red gate. **(C)** Percentage of mCherry-positive alveolar macrophages out of total alveolar macrophages as determined by flow cytometry from n=6 mice administered ARMMs.mCherry.P. **(D)** Paraffin-embedded lung sections were immuno-stained for CD206 (macrophage marker) and mCherry to visualize colocalization. mCherry signal was observed in macrophages in lung cross sections from mice administered with ARMMs.mCherry.P but not PBS. (**E**) Biodistribution time course of ARMMs.mCherry.P in lung. Lung cross-sections displaying mCherry signal (red) with nuclear counterstain (blue) at multiple timepoints after oropharyngeal aspiration of ARMMs.mCherry.P. Scale bar 100um.

**Supplementary Fig.S9.**
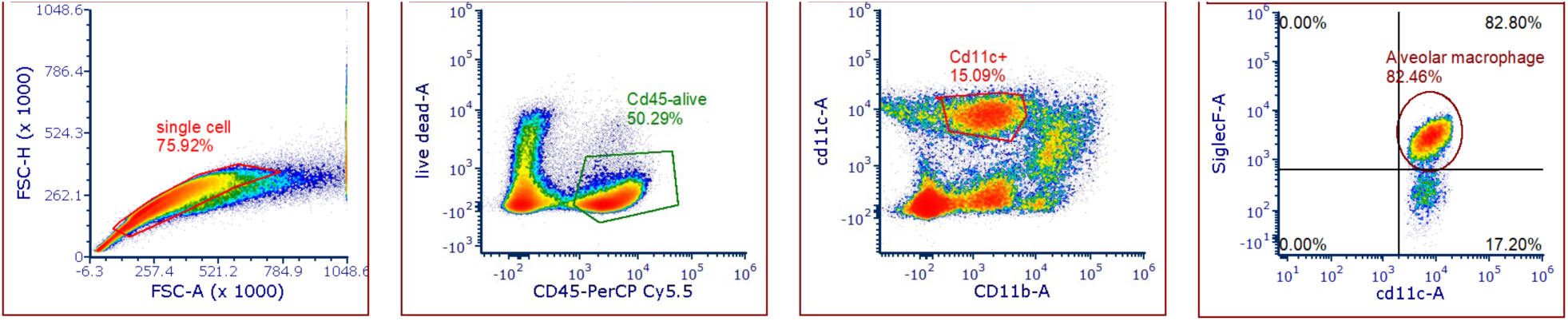
Gating strategy for alveolar macrophages. Full lung digestion (see methods) was followed by flow cytometry analysis. Alveolar macrophages are defined as CD45^+^ CD11c^hi^ CD11b^mid^ SiglecF^+^ cells. Cells were first gated on FSC-A vs FSC-H to exclude doublets, then CD45+ live cells were gated, followed by gating of CD11c^hi^ CD11b^mid^ cells, and finally by gating SiglecF^+^ cells.

**Supplementary Fig.S10.**
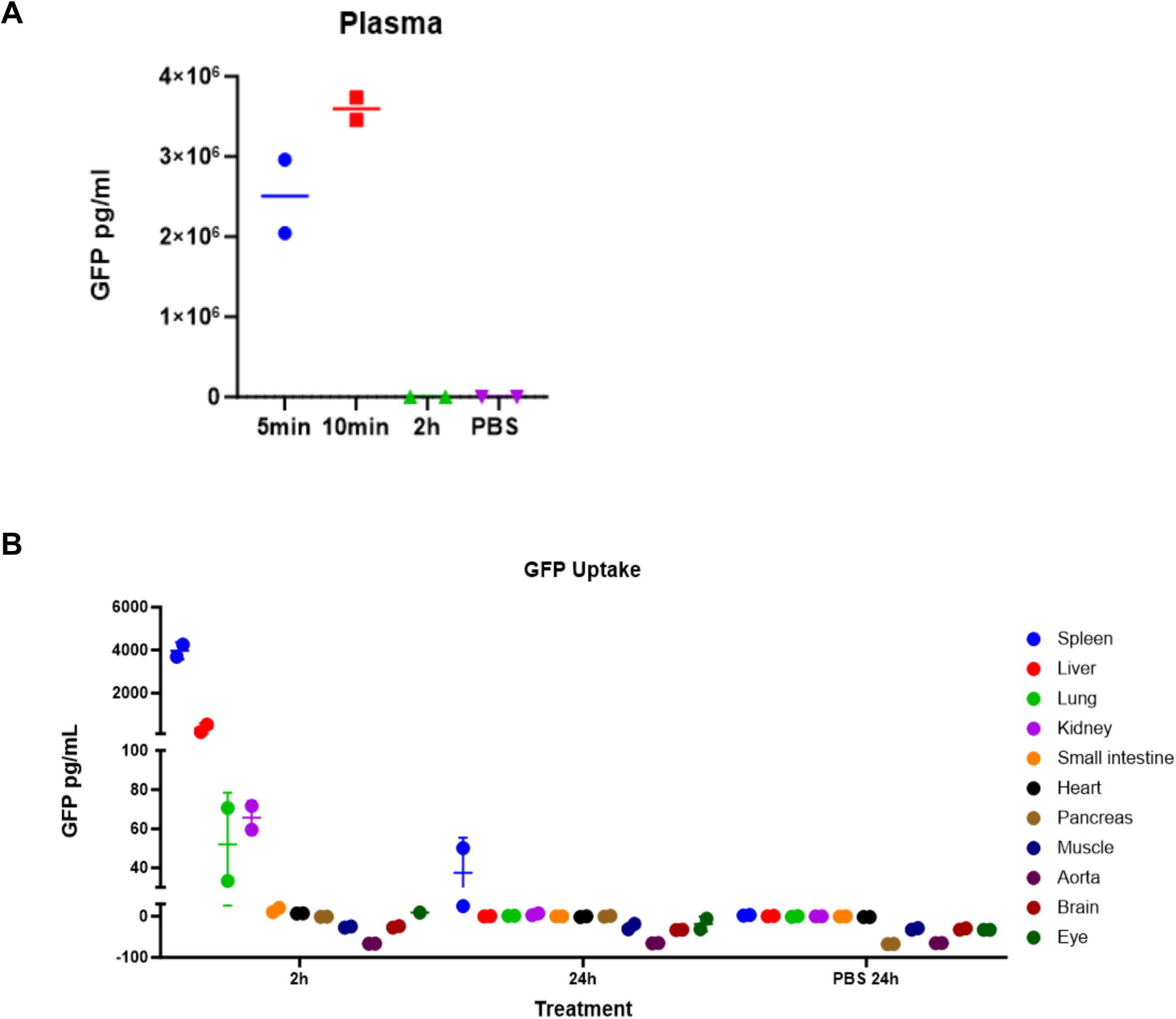
*In vivo* biodistribution of ARMMs.GFP.P by retro-orbital intravenous (IV) injection in mice. 1×10^12^ of ARMMs were injected for the *in vivo* biodistribution studies with two animals per group. GFP signals were measured by ELISA and normalized by 300ug of protein for tissues or 50uL of plasma. (**A**) Pharmacokinetic evaluation of ARMMs in plasma. (**B**) Time course measurement of GFP in various tissues with significant uptake of ARMMs.

**Supplementary Fig.S11.**
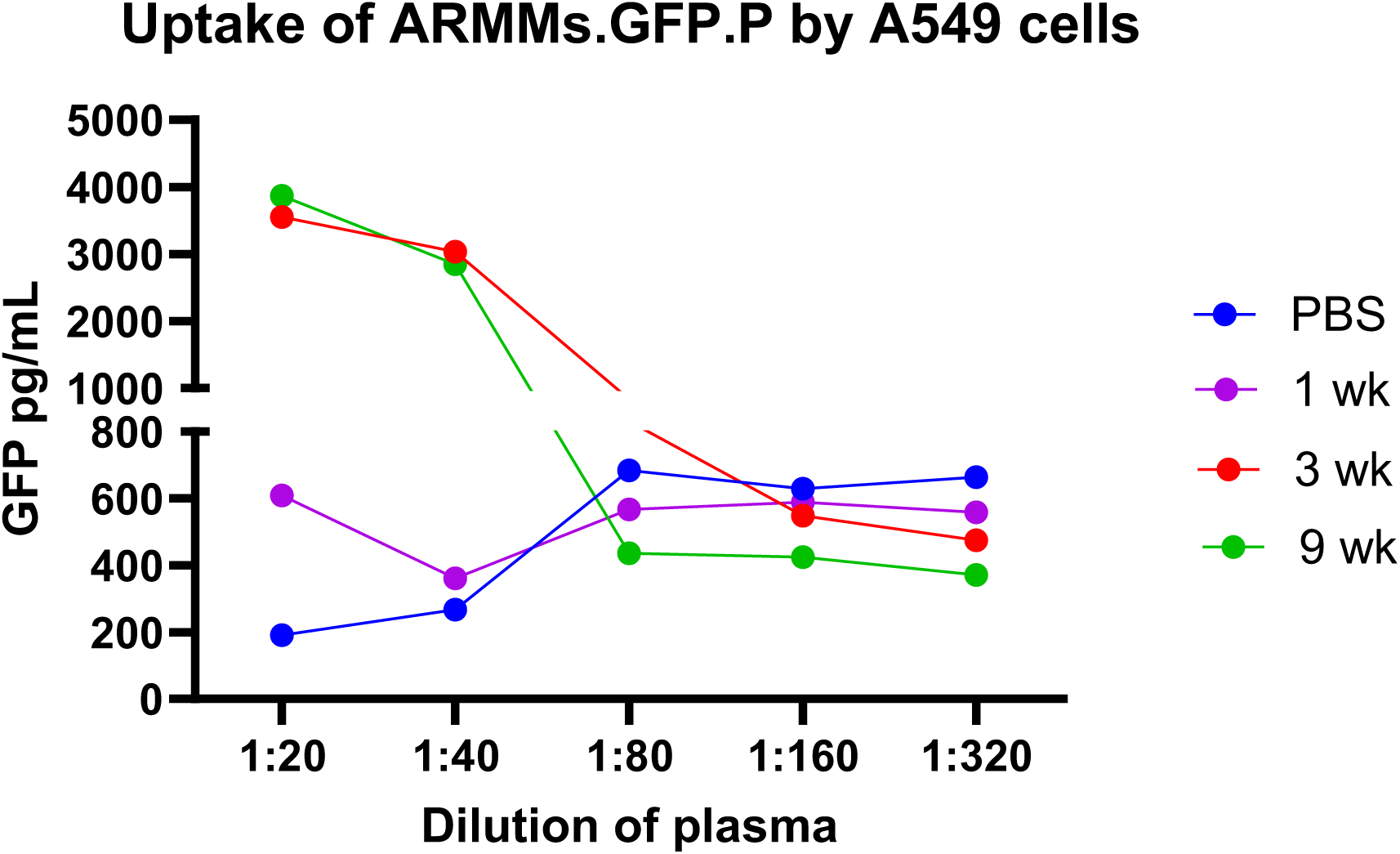
Intravenous administration of ARMMs does not result in generation of neutralizing antibodies. Mouse plasma were collected at day 28 from mice injected with ARMMs.GFP.P with either1 dose or 3 doses weekly of 1×10^12^ ARMMs per injection. The presence of neutralizing antibodies in plasma was measured by blockade of ARMMs.GFP.P uptake. A549 cells were treated with ARMMs.GFP.P at 1×10^5^ particles/cell for 4h. After 3x washes, GFP was quantified from cells by ELISA. No inhibition of ARMMs.GFP.P uptake by A549 cells was observed.

**Supplementary Fig.S12.**
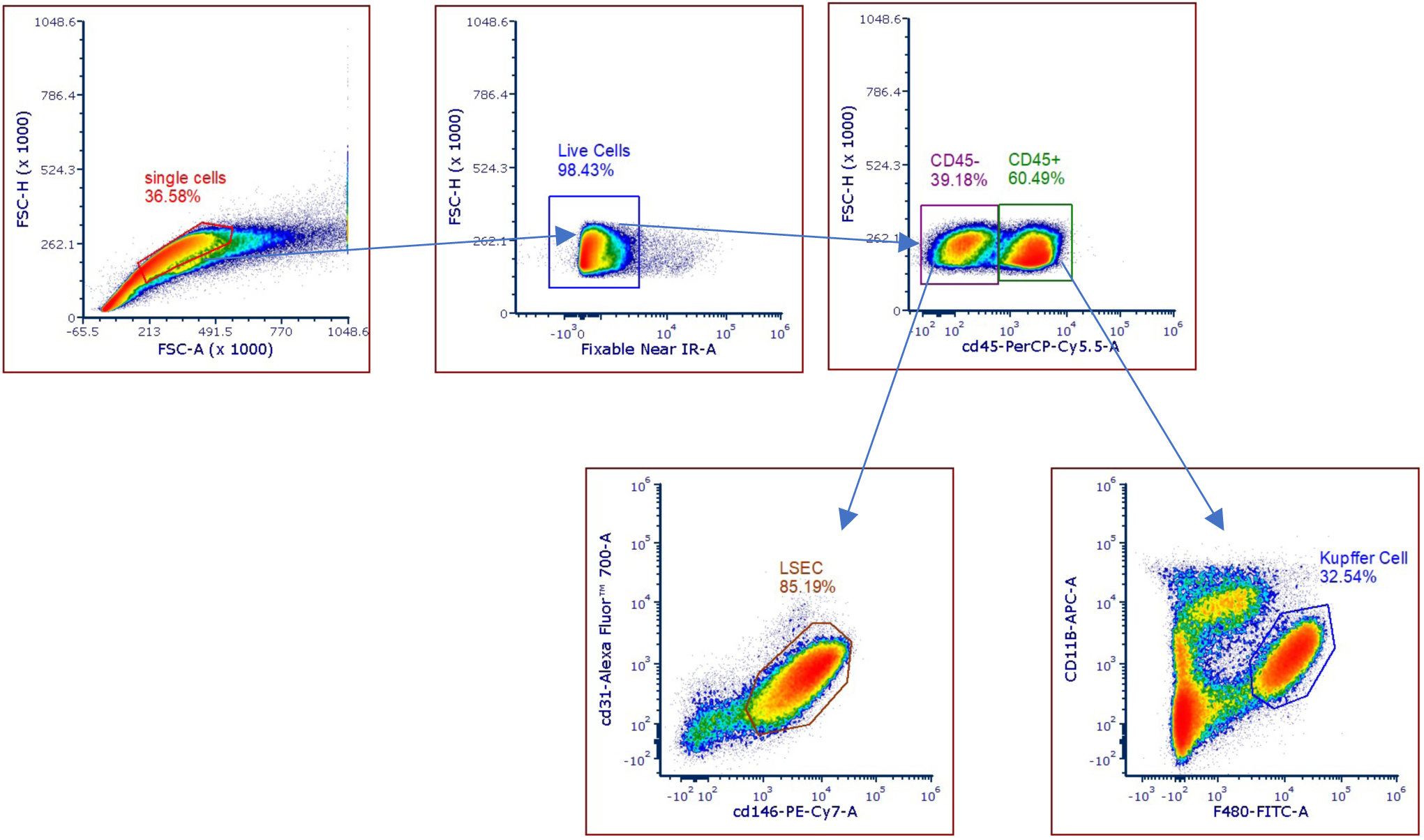
Liver cell gating strategy for LSEC and KC. Non-parenchymal cells were first gated based on FSC-A vs FSC-H to remove doublets. Dead cells were then excluded using Live/Dead fixable near IR stain. CD45^-^ cells were gated to allow for identification of LSEC based on expression of CD31^+^ and CD146^+^ markers. CD45^+^ cells were gated and Kupffer cells were identified based on F4/80^+^ and CD11b^mid^ markers.

**Supplementary Fig.S13.**
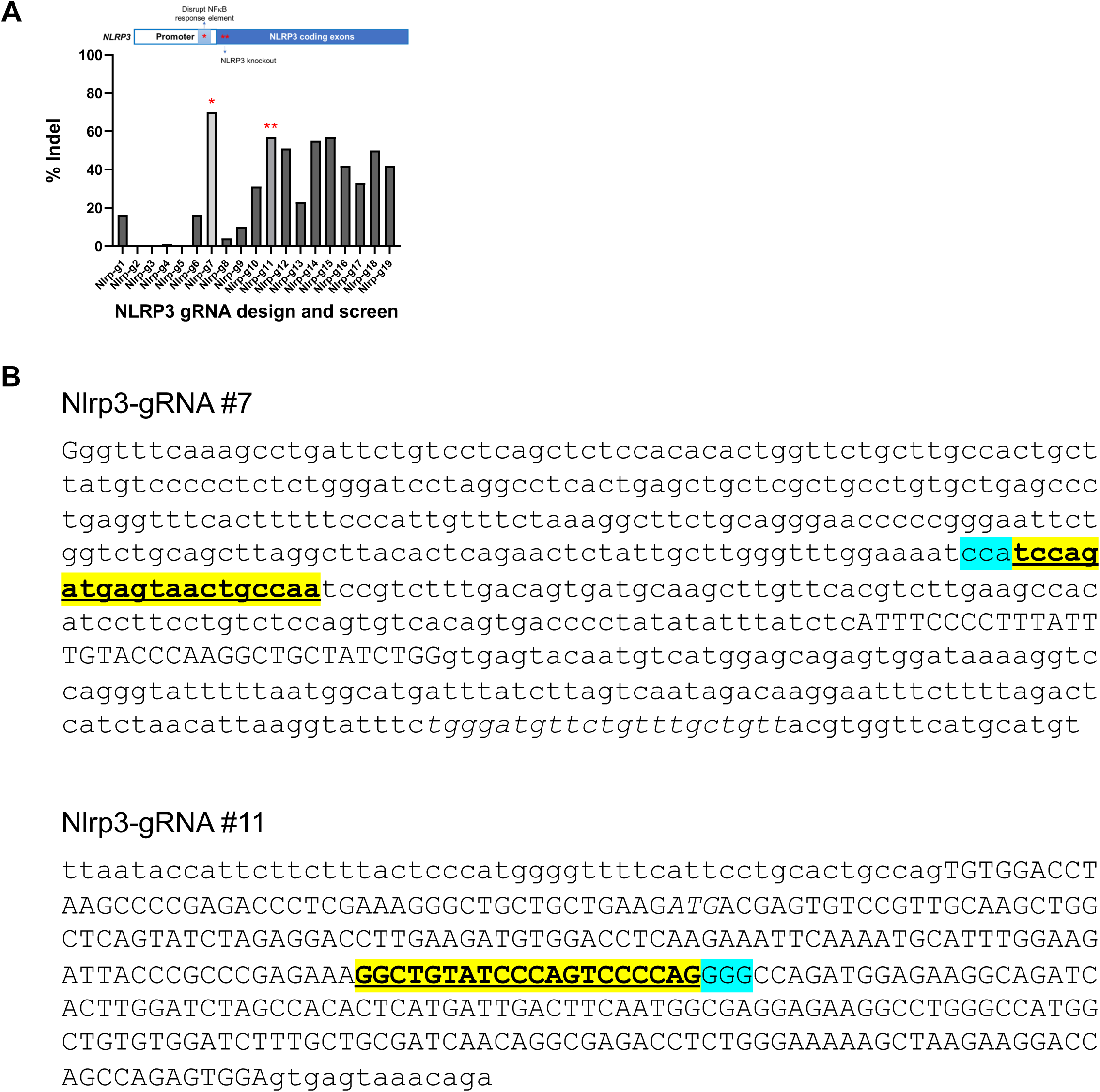
Guide RNA screen for targeting the mouse *Nlrp3* gene. (**A**) NGS data showing editing efficiency of the mouse Nlrp3 gene in mouse Neuro2a cells by transient transfection of plasmids expressing ARRDC1-Cas9 fusion protein and Nlrp3 targeting gRNAs. (**B**) Nlrp3 gRNA sequences that yielded highest level of editing. Nlrp3-gRNA #7 in reverse direction is yellow highlighted in the mouse Nlrp3 promoter region. Nlrp3-gRNA #11 in the forward direction is yellow highlighted in the first exon. PAM sequences are highlighted in blue.

**Supplementary Fig.S14.**
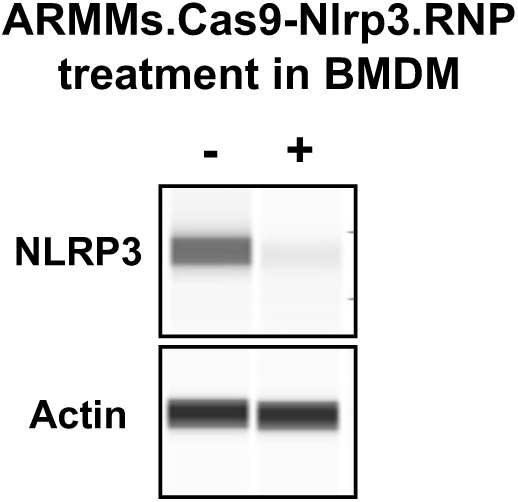
Reduced NLRP3 protein in bone marrow-derived macrophages after editing by ARMMs.Cas9-Nlrp3^#11^.RNP. Western blot data showing protein expression levels of mouse Nlrp3 in BMDMs after ARMMs treatment. β-actin was used as lane loading control.

**Supplementary Fig.S15.**
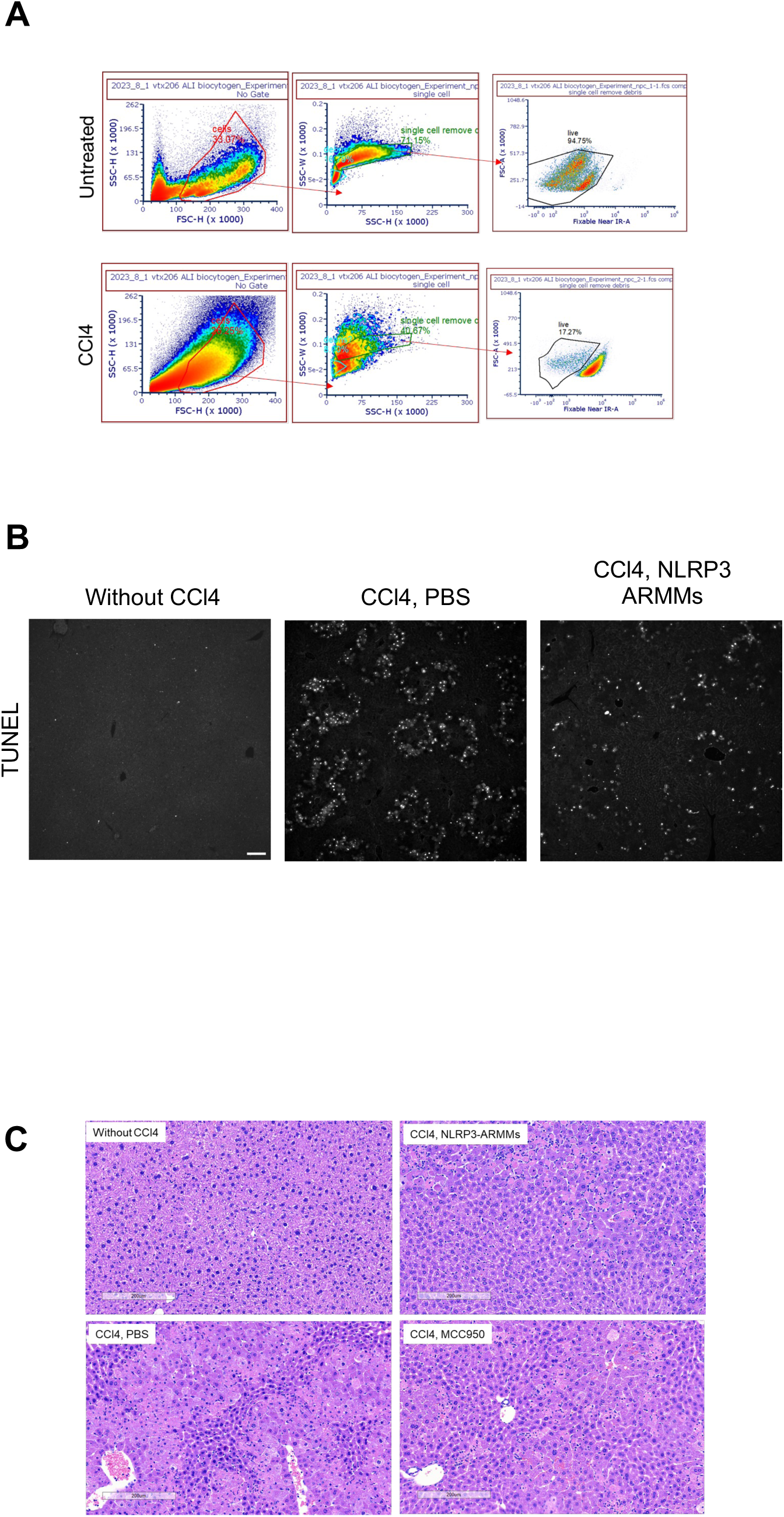
**(A)** Representative density plots depicting the gating strategy used to evaluate percentage of live cells from single-cell suspensions of livers from untreated or CCl4-treated mice. Livers from mice were homogenized as described in the methods section, and single cell suspensions were first gated based on FSC-H and SSC-H to remove debris, and further gated on SSC-H and SSC-W to select for single cell-events. Cells were finally gated based on live-dead signal. **(B)** Representative images of TUNEL stain from paraffin-embedded liver (left lobe) cross sections. Sections were evaluated by TUNEL assay as described in the methods section, and five fields of view were captured for each section encompassing a large portion of the tissue. Representative TUNEL-stained images are shown with examples from CCl4-treated mice administered either PBS or Cas9-gRNA RNPs targeting NLRP3. Mice not treated with CCl4 displayed no TUNEL signal. Scale bar 100 µm **(C)** Left lobe cross-sections were stained for H&E to evaluate tissue necrosis. Representative images for each condition are shown.

## References

1 Bulaklak, K. & Gersbach, C. A. The once and future gene therapy. Nature Communications 11 (2020). 10.1038/s41467-020-19505-2

2 Ling, Q., Herstine, J. A., Bradbury, A. & Gray, S. J. AAV-based in vivo gene therapy for neurological disorders. Nature Reviews Drug Discovery 22, 789–806 (2023). 10.1038/s41573-023-00766-7

3 Hou, X., Zaks, T., Langer, R. & Dong, Y. Lipid nanoparticles for mRNA delivery. Nature Reviews Materials 6, 1078–1094 (2021). 10.1038/s41578-021-00358-0

4 Ertl, H. C. J. Immunogenicity and toxicity of AAV gene therapy. Frontiers in Immunology 13 (2022). 10.3389/fimmu.2022.975803

5 Guo, C., Ma, X., Gao, F. & Guo, Y. Off-target effects in CRISPR/Cas9 gene editing. Frontiers in Bioengineering and Biotechnology 11 (2023). 10.3389/fbioe.2023.1143157

6 Hanlon, K. S. et al. High levels of AAV vector integration into CRISPR-induced DNA breaks. Nature Communications 10 (2019). 10.1038/s41467-019-12449-2

7 Kim, D., Luk, K., Wolfe, S. A. & Kim, J.-S. Evaluating and Enhancing Target Specificity of Gene-Editing Nucleases and Deaminases. Annual Review of Biochemistry 88, 191–220 (2019). 10.1146/annurev-biochem-013118-111730

8 Jiang, Z. & Dalby, P. A. Challenges in scaling up AAV-based gene therapy manufacturing. Trends in Biotechnology 41, 1268–1281 (2023). 10.1016/j.tibtech.2023.04.002

9 Youssef, M., Hitti, C., Puppin Chaves Fulber, J. & Kamen, A. A. Enabling mRNA Therapeutics: Current Landscape and Challenges in Manufacturing. Biomolecules 13, 1497 (2023). 10.3390/biom13101497

10 Nabhan, J. F., Hu, R., Oh, R. S., Cohen, S. N. & Lu, Q. Formation and release of arrestin domain-containing protein 1-mediated microvesicles (ARMMs) at plasma membrane by recruitment of TSG101 protein. Proc Natl Acad Sci U S A 109, 4146–4151 (2012). 10.1073/pnas.1200448109

11 Wang, Q. et al. ARMMs as a versatile platform for intracellular delivery of macromolecules. Nat Commun 9, 960 (2018). 10.1038/s41467-018-03390-x

12 Jeppesen, D. K., Zhang, Q., Franklin, J. L. & Coffey, R. J. Extracellular vesicles and nanoparticles: emerging complexities. Trends in Cell Biology 33, 667–681 (2023). 10.1016/j.tcb.2023.01.002

13 Fiumara, M. et al. Genotoxic effects of base and prime editing in human hematopoietic stem cells. Nature Biotechnology (2023). 10.1038/s41587-023-01915-4

14 Leibowitz, M. L. et al. Chromothripsis as an on-target consequence of CRISPR–Cas9 genome editing. Nature Genetics 53, 895–905 (2021). 10.1038/s41588-021-00838-7

15 Cullot, G. et al. CRISPR-Cas9 genome editing induces megabase-scale chromosomal truncations. Nature Communications 10 (2019). 10.1038/s41467-019-09006-2

16 Luther, K. et al. Scalable Production and Purification of Engineered ARRDC1-Mediated Microvesicles in a HEK293 Suspension Cell System (Cold Spring Harbor Laboratory, 2024).

17 Lee, Y., Jeong, M., Park, J., Jung, H. & Lee, H. Immunogenicity of lipid nanoparticles and its impact on the efficacy of mRNA vaccines and therapeutics. Experimental & Molecular Medicine 55, 2085–2096 (2023). 10.1038/s12276-023-01086-x

18 Chen, S. P. & Blakney, A. K. Immune response to the components of lipid nanoparticles for ribonucleic acid therapeutics. Current Opinion in Biotechnology 85, 103049 (2024). 10.1016/j.copbio.2023.103049

19 Alameh, M.-G. et al. Lipid nanoparticles enhance the efficacy of mRNA and protein subunit vaccines by inducing robust T follicular helper cell and humoral responses. Immunity 54, 2877–2892.e2877 (2021). 10.1016/j.immuni.2021.11.001

20 Jeppesen, D. K. et al. Reassessment of Exosome Composition. Cell 177, 428–445.e418 (2019). 10.1016/j.cell.2019.02.029

21 Madisen, L. et al. A robust and high-throughput Cre reporting and characterization system for the whole mouse brain. Nature Neuroscience 13, 133–140 (2010). 10.1038/nn.2467

22 Tsai, S. Q. et al. GUIDE-seq enables genome-wide profiling of off-target cleavage by CRISPR-Cas nucleases. Nature Biotechnology 33, 187–197 (2015). 10.1038/nbt.3117

23 Davis, J. R., et al. Efficient in vivo base editing via single adeno-associated viruses with size-optimized genomes encoding compact adenine base editors. Nature Biomedical Engineering 6, 1272–1283 (2022). 10.1038/s41551-022-00911-4

24 Rothgangl, T. et al. In vivo adenine base editing of PCSK9 in macaques reduces LDL cholesterol levels. Nature Biotechnology 39, 949–957 (2021). 10.1038/s41587-021-00933-4

25 Huang, X. et al. The landscape of mRNA nanomedicine. Nature Medicine 28, 2273–2287 (2022). 10.1038/s41591-022-02061-1

26 Blevins, H. M., Xu, Y., Biby, S. & Zhang, S. The NLRP3 Inflammasome Pathway: A Review of Mechanisms and Inhibitors for the Treatment of Inflammatory Diseases. Frontiers in Aging Neuroscience 14 (2022). 10.3389/fnagi.2022.879021

27 Almuttaqi, H. & Udalova, I. A. Advances and challenges in targeting IRF5, a key regulator of inflammation. The FEBS Journal 286, 1624–1637 (2019). 10.1111/febs.14654

28 Scholten, D., Trebicka, J., Liedtke, C. & Weiskirchen, R. The carbon tetrachloride model in mice. Laboratory Animals 49, 4–11 (2015). 10.1177/0023677215571192

29 Liu, Y. et al. S100A8-Mediated NLRP3 Inflammasome-Dependent Pyroptosis in Macrophages Facilitates Liver Fibrosis Progression. Cells 11, 3579 (2022). 10.3390/cells11223579

30 Sharma, P., Hoorn, D., Aitha, A., Breier, D. & Peer, D. The immunostimulatory nature of mRNA lipid nanoparticles. Advanced Drug Delivery Reviews 205, 115175 (2024). 10.1016/j.addr.2023.115175

31 Nakano, H., Nakano, K. & Cook, D. N. Isolation and Purification of Epithelial and Endothelial Cells from Mouse Lung. Methods Mol Biol 1799, 59–69 (2018). 10.1007/978-1-4939-7896-0_6

32 Conant, D. et al. Inference of CRISPR Edits from Sanger Trace Data. CRISPR J 5, 123–130 (2022). 10.1089/crispr.2021.0113

33 Clement, K. et al. CRISPResso2 provides accurate and rapid genome editing sequence analysis. Nat Biotechnol 37, 224–226 (2019). 10.1038/s41587-019-0032-3

34 Doman, J. L., Raguram, A., Newby, G. A. & Liu, D. R. Evaluation and minimization of Cas9-independent off-target DNA editing by cytosine base editors. Nat Biotechnol 38, 620–628 (2020). 10.1038/s41587-020-0414-6

35 Concordet, J. P. & Haeussler, M. CRISPOR: intuitive guide selection for CRISPR/Cas9 genome editing experiments and screens. Nucleic Acids Res 46, W242–W245 (2018). 10.1093/nar/gky354

36 Hsu, P. D. et al. DNA targeting specificity of RNA-guided Cas9 nucleases. Nat Biotechnol 31, 827–832 (2013). 10.1038/nbt.2647

37 Tycko, J. et al. Mitigation of off-target toxicity in CRISPR-Cas9 screens for essential non-coding elements. Nat Commun 10, 4063 (2019). 10.1038/s41467-019-11955-7

38 Doench, J. G. et al. Optimized sgRNA design to maximize activity and minimize off-target effects of CRISPR-Cas9. Nat Biotechnol 34, 184–191 (2016). 10.1038/nbt.3437

39 Moreno-Mateos, M. A. et al. CRISPRscan: designing highly efficient sgRNAs for CRISPR-Cas9 targeting in vivo. Nat Methods 12, 982–988 (2015). 10.1038/nmeth.3543

